# Elucidating Odorant Masking Effects in Food Matrices Through Olfactory Receptor-Based Profiling

**DOI:** 10.64898/2026.01.13.699200

**Authors:** Boyong Hu, Haochen Zheng, Hong Liang, Haoming Lv, Qing Xiao, Yuqin Jiang, Qi Lu, Yunwei Niu, Heng Wang, Zuobing Xiao

## Abstract

Dynamic interactions among aroma compounds, especially odorant masking, constitute a key determinant of the intricate flavor profiles of thermally processed foods. However, the molecular and receptor-level mechanisms governing mutual odor suppression remain largely unclear. This study investigated the reciprocal sensory masking between 2,3,5-trimethylpyrazine and L-menthol via systematic sensory evaluation. Olfactory receptor screening identified OR5K1 as the responsive receptor for 2,3,5-trimethylpyrazine (EC_50_ = 27.67 ± 2.02 μmol/L) and OR2W1 as the functional receptor for L-menthol (EC_50_ = 22.69 ± 3.85 μmol/L). Consistently, quantitative cellular response assays further validated that 2,3,5-trimethylpyrazine and L-menthol exert mutual inhibitory effects on their respective receptors, as evidenced by the significant inhibition of receptor signaling. Moreover, molecular modeling predicted that 2,3,5-trimethylpyrazine and L-menthol competitively occupy the same orthosteric pocket via analogous binding conformations. These results provide novel receptor-level insights into odorant inhibitory effects, connecting sensory perception to molecular recognition, and deepen mechanistic understanding of flavor perception.

## 1 INTRODUCTION

Food flavors exhibit remarkable diversity resulting from complex interactions among odorant molecules. These interactions primarily manifest through flavor enhancement (intensification of specific aroma notes) and flavor masking (suppression of specific aroma notes). Masking effects are particularly significant in food science ^1^, enabling targeted modification of undesirable flavors while preserving desirable sensory profiles. Practical strategies include molecular encapsulation, enzymatic modification, and strategic blending with masking compounds ^1–3^. Notably, aroma-induced masking allows flavor modulation without altering food matrices, offering broad applications in food development. Aroma masking is ubiquitous, exemplified by spice-derived compounds suppressing baked/burnt odors in thermally processed foods ^4, 5^. A representative system involves interactions between Maillard-derived 2,3,5-trimethylpyrazine (TMP, imparting nutty-roasted notes) and spice-originated L-menthol (L-MT, providing herbal/minty aromas) ^6–8^. Their co-occurrence in grilled meats and baked goods provides an ideal model for studying multisensory integration and olfactory variations. Furthermore, the masking effects among aroma compounds are strongly modulated by complex food matrices. Food components such as lipids, proteins, polysaccharides and salts can bind, partition or sequester volatile compounds, thereby changing their release behavior, perceived intensity and mutual interactions ^9^. Such matrix effects directly regulate the masking mechanisms at olfactory receptor and perceptual levels. The occurrence and intensity of aroma masking in real food systems depend on both the intrinsic interactions among aroma compounds and the extrinsic modulation of the food matrix, which collectively shape the overall aroma profile of food products.

The multifaceted nature of food flavors arises from a dynamic interplay between odorant molecules, with synergistic intensification and competitive masking of specific aroma notes being principal manifestations. The investigation of aroma interactions in foods requires an integrated approach combining instrumental analysis with sensory evaluation. Gas chromatography-olfactometry-mass spectrometry (GC-O-MS) enables precise identification of key odorants and their interactions in complex matrices such as coffee, wine, and processed meats ^10, 11^, while sensory methods like S-curve modeling quantitatively characterize flavor thresholds and synergistic/antagonistic relationships ^12^. As evidenced by the studies of Li et al. and Guo et al., acetate significantly enhances floral-fruity notes in wines while aldehydes demonstrate sweet-enhancing effects in oranges through cross-modal olfactory-gustatory interactions ^13, 14^. These diverse methodologies collectively bridge chemical composition with perceptual outcomes, thereby providing deeper insights into flavor modulation mechanisms essential for food quality optimization. Recognizing the inherent limitations of sensory evaluation’s subjectivity, contemporary research increasingly incorporates molecular biological approaches to unravel the fundamental mechanisms underlying odorant interactions. Olfactory perception is the biological process of detecting, identifying, and interpreting volatile chemical substances in the environment through the olfactory system. Crucially, this process relies on olfactory receptors (ORs, ≈ 400 types in humans) located in the olfactory epithelium of the upper nasal cavity, which specifically bind to particular types of odor molecules.

Comprehensive mapping of olfactory receptor activation profiles has revolutionized our understanding of flavor perception, particularly through the elucidation of precise OR-odorant binding specificities ^15, 16^. For instance, researchers identified OR2T11 as a small-molecule thiol-responsive receptor and uncovering the crucial role of metal ions in mediating odorant-receptor interactions ^17^. After deorphanizing olfactory receptors and identifying ligand-receptor pairs, in-depth analyses of polymorphisms in olfactory receptor coding genes have been further conducted. For example, genetic variations in OR7D4 influence individual differences in the valence (pleasantness or unpleasantness) and intensity of perceived steroid odors such as androstenone ^18, 19^. *In vitro* heterologous expression systems of olfactory receptors can also be used to study the interactions between aroma compounds. For example, the synergistic effect of aldehydes on muscone can be investigated using its corresponding receptor OR5AN1 ^20–23^. Moreover, studies on molecular modeling, molecular dynamics, and meta-databases of olfactory receptor-odorant interactions have contributed to the understanding of olfactory receptor activation mechanisms and the discovery of potential odorant receptors ^24, 25^. The prediction of active pockets and binding sites also further advances research on the structural and functional properties of receptor proteins at the microscopic level.

In this study, the olfactory detection thresholds of 2,3,5-trimethylpyrazine and L-menthol, and olfactory perceptual interactions of 2,3,5-trimethylpyrazine with L-menthol were determined through sensory experiments. Subsequently, ORs heterologous expression model cells were constructed and *in vitro* cellular experiments were carried out to screen for 2,3,5-trimethylpyrazine-responsive receptors. OR5K1 was shown to be a highly responsive receptor for 2,3,5-trimethylpyrazine and the effects of L-menthol on the 2,3,5-trimethylpyrazine-OR5K1 pair were explored. Additionally, as a receptor that can be activated by L-menthol, OR2W1 was used to detect the modulation of L-menthol receptor activation by 2,3,5-trimethylpyrazine. Through analyzing the modulation effects of the two odorants on their corresponding receptors, their interaction patterns were revealed. Finally, molecular modeling was used to explain the potential mechanism of odorants’ interaction at the molecular level.

## 2 MATERIALS AND METHODS

### 2.1 Chemicals

Chemicals required for the cellular experiments: DMEM medium, fetal bovine serum (FBS) (Premium Plus), 1×PBS buffer, 10000 U/mL penicillin/streptomycin, trypsin/EDTA solution, 1mol/L HEPES, 1×HBSS, and Opti-MEM medium were purchased from Invitrogen (California, America). Lipofectamine™ 3000 purchased from Thermo Fisher Scientific (Shanghai, China) was used for cell transfection. The reagents used for odorant stimulation include 2,3,5-trimethylpyrazine and L-menthol, all with a purity of >99%, sourced from Aladdin (Shanghai, China). The odorant solvent is dimethyl sulfoxide and dipropylene glycol (Beyotime, Shanghai, China).

### 2.2 Sensory conditions and sensory panel

All sensory evaluation procedures were conducted in strict compliance with relevant regulatory guidelines (ISO 8586:2012; ISO 8589:2007) and institutional ethical standards. The sensory team consisted of 20 trained members, including 10 males and 10 females. Participants must be in good health, free from any conditions that may impair olfactory function. During the training process, panelists are required to undergo olfactory threshold and identification tests to ensure olfactory sensitivity and exclude individuals with olfactory abnormalities. Furthermore, panelists receive repeated training in odor recognition and description, intensity quantification, and difference discrimination to reinforce their perception and memory of the characteristics of each experimental odor. Following 60 hours of pre-experimental training (2 hours daily), only qualified members were selected to participate in the formal trials ^26, 27^.

All experimental samples were prepared in full compliance with national food safety regulations, with tested concentrations maintained within safety thresholds to ensure human health. The sensory evaluation protocol has been approved by the Ethics Committee of Shanghai Jiao Tong University (Approval Number: B20250330I) and all participants have been informed of the experimental purpose, protocol, and ethical review outcome.

### 2.3 Sensory experiments to verify odor interaction

A systematic concentration-gradient approach was implemented to quantify odor interaction between target aromatics, following our established protocols for binary mixture analysis ^14, 28^. The thresholds of aroma compounds were determined by the three-alternative forced-choice method, adhering to the ISO 13301:2018 (***Sensory analysis-Methodology-General guidance for measuring odor, flavor and taste detection thresholds by a three-alternative forced-choice (3-AFC) procedure***). The experiment on the interaction between 2,3,5-trimethylpyrazine and L-menthol was conducted with modifications based on the method reported by Trimmer et al. (see Supporting Information for details) ^23^. Specifically, the odorants at the required concentration gradients were diluted with dipropylene glycol (DiPG) to prepare liquid solutions, which were then transferred to circular cellulose filter papers (2 cm in diameter). The filter papers were immersed in the solutions and allowed to absorb the odorants for 10 minutes until saturation. Subsequently, the odorant-loaded filter papers were placed in 25 mL brown glass vials with lids and left at room temperature for 2.5 hours to ensure the sufficient volatilization and equilibration of the odorants on the papers prior to the olfactory tests. All olfactory tests were required to be completed within 4 hours.

The experimental data were analyzed to determine olfactory thresholds and odorant interactions using the S-curve method, with the following fitting equation (formula 1). Here, *P* represents the chance-corrected probability of detection, *x* is the concentration of the odorant, *c* is the olfactory threshold, and *D* is a parameter defining the slope of the function ^29^:

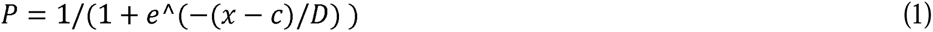

For binary odorant systems, two odorants, A and B were mixed, and the evaluators sniffed the mixture to detect odorant A. The measured probability of detecting A, denoted as P(A) was compared to the theoretical probability of detection, p(A), for A alone. The lg(c)-p curve was plotted and fitted, where the concentration values were summed while maintaining the original proportion of A and B. The threshold was defined as the concentration corresponding to P=0.5 on the fitted curve. If P(A) at P=0.5 was lower than p(A), odorant B exhibited a masking effect on A (S-curve right shift). If P(A) was higher than p(A), B demonstrated a mutual enhancement or synergistic effect on A (S-curve left shift). If P(A) was equal to p(A), no significant interaction between A and B was observed ^29^.

### 2.4 ORs plasmids preparation

The encoding sequences of ORs were amplified from human genomic DNA via polymerase chain reaction (PCR) using specific primers and subsequently ligated into the pCI mammalian expression vector (Promega, Shanghai) using T4-DNA ligase (Fig. S1). The sequence information for each gene is available in GenBank (http://www.ncbi.nlm.nih.gov/genbank/). Similarly, 294 ORs (haplotype, excluding variants) were cloned and ligated to expression plasmids for receptor screening (Table S1; Table S2). Plasmid sequencing was performed using Oxford Nanopore Technology (Cowin Biotech, Shanghai, China).

### 2.5 Cell culture and construction of ORs expression system

Hana-3A cells, an engineered derivative of the human embryonic kidney cell line (HEK293), were used for olfactory receptor expression ^20^. Hana-3A cells were cultured with DMEM (with 4.5 g/L D-glucose) supplemented with 10% FBS, 1× penicillin-streptomycin (100 U/mL), and 2 mmol/L L-glutamine in 10 cm dishes, and incubated at 37°C, 5% COLJ, and 100% humidity. After cell confluence reaches 100%, perform passaging and continue seeding cells in 10 cm dishes and then transfected via lipofection when reached 60% confluence. 8000 ng of the corresponding ORs and pGloSensor-22F plasmid (Promega, Shanghai), and 4000 ng of mRTP1s and Gα_olf_ plasmid were co-transfected ^30^.

### 2.6 Odorants preparation

All odorants are diluted from the pure reagents by dimethyl sulfoxide (DMSO). Stock solutions of each odorant were first prepared at a concentration of 2 mol/L and then sequentially diluted to suitable concentrations. The odorants required for the interaction experiments were formulated in advance, followed by premixing of the stock solutions in the desired proportions prior to dilution for use as operating solutions. This approach ensures that both single-component and mixed odorants are administered at a consistent volume of 1 μL, minimizing potential variability introduced during the delivery process. Gas chromatography-quadrupole/orbitrap mass spectrometry (GC-Q/Orbitrap MS, Thermo Fisher Scientific, America) was used to analyze the components of the pre-configured binary systems.

### 2.7 Cell viability assay

The methylthiazolyldiphenyl-tetrazolium bromide viability assay (MTT viability assay) was employed to assess the cytotoxicity of the applied odorants ^31^. Cell viability was measured after transfection and subsequent odorant application, with comparisons made between the odorant-treated group and both the blank group (untreated) and solvent control (DMSO) group.

### 2.8 Luminescence assay

Cells after 24 h of transfection were resuspended with 87% HBSS, 10% FBS, and 3% GloSensor cAMP Reagent (Promega, Wisconsin, America) and plated in a white 96-well plate (Thermo Scientific™ Nunc™ F96 MicroWell™) ^32^. For the luminescence assay, the microplate reader (Thermo Fisher Varioskan LUX) and Skanlt Software 6.1 RE (ver.6.1.0.51) were used. After incubating for 2 h at room temperature in the dark, the baseline of each well was recorded preferentially, and then prepared operating solutions were added to the wells to obtain suitable final concentrations on the cells. Immediately thereafter, the luminescence signal was determined and kinetic cycle testing was used so that the luminescence signal in each well was detected at 30s intervals ^22^.

### 2.9 Analysis of cAMP luminescence signals

All data points of the baseline and data points after odorant application were averaged separately. The mock control (solvents for odorants) was subtracted from the luminescence value of each well and divided by the basal level, and the result expressed in multiplicity as a dimensionless constant was taken as the luminescence intensity (formula 2).

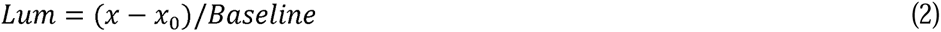

The dataset was normalized to the maximum amplitude of the reference odorant-receptor pair ^33^, and the concentration-response curves were fitted using the Logistic function in Origin 2019 software. All data are expressed as mean ± SD and error bars are indicated on the data points of all the figures.

### 2.10 Molecular modeling

#### 2.10.1 Molecular docking

AlphaFold3 supports the de novo prediction of complex structures from sequence alone, eliminating the need for a predefined receptor structure or binding pocket. In contrast to conventional docking approaches, it jointly models the complete 3D conformations of the protein and ligand at the same time, enabling accurate characterization of their mutual structural adaptation. AlphaFold3 1.0 was employed to predict the interactions between aroma compounds and ORs. Mature protein sequence of OR5K1 (UniProt ID: Q8NHB7) and OR2W1 (UniProt ID: Q9Y3N9), and two aroma compounds (2,3,5-trimethylpyrazine and L-menthol) were selected to investigate their binding pattern. Protein sequences of ORs and simplified molecular input line entry system (SMILES) of compounds served as an input to AlphaFold3. Each ligand-receptor pair was individually predicted and a total of complex structures were obtained. Intermolecular interactions of each complex were analyzed using Maestro Viewer 14.3, with results presented as a 2D interaction diagram. Additionally, 3D interaction visualizations were created using PyMOL 2.3.0.

#### 2.10.2 Molecular dynamics simulation

The initial conformations of the complexes were predicted using AlphaFold3. The systems for simulation were built via CHARMM-GUI website, where the bilayer was composed of ∼ 169 1-palmitoyl-2-oleoyl-sn-glycero-3-phosphocholine (POPC) ^34^. The bilayer and protein complex were solvated into periodic hexagonal type TIP3P water box containing 0.15 M KCl. Molecular dynamics simulations were performed using GROMACS 2020, with the CHARMM36m force field for proteins and the CHARMM General Force Field (CGenFF) for ligands ^35^. Energy minimization and equilibration utilized the default parameters generated by the CHARMMGUI webserver. In the production, the systems were simulated for 200 ns with a 2-fs time step. During the simulation, the vLJrescale thermostat was used to keep the temperature at 303.15 K and the ParrinelloLJRahman semi-isotropic barostat controlled pressure at 1LJatm (NPT ensemble) ^36^. Van der Waals interactions were force-switched off from 0.8 to 1.2 nm. The Particle Mesh Ewald method was used with the 1.2 nm cutoff ^37^. The LINCS algorithm was applied to constrain all bonds associated with hydrogen atoms ^38^. The root mean square deviation (RMSD) of the ligand heavy atoms was calculated using the GROMACSLJrms tool, with the predicted initial conformation as the reference. The binding free energy was calculated using the molecular mechanics/Poisson Boltzmann surface area (MM/PBSA) method ^39^. In this work, the change in conformational entropy is omitted owing to its high computational cost and limited predictive accuracy. Binding free energies for all protein–ligand complexes were calculated using gmx_MMPBSA 1.6.2 ^40^, based on 200LJns (2000LJframe) molecular dynamics trajectories.

### 2.11 Data analysis

All of the experiments were performed in appropriate biological replicates and the results of statistical analysis were presented as mean ± SEM (standard error of mean). Microsoft Office Excel was used for data processing and GraphPad Prism 10 and Origin 2019 software were used to graph the data. For comparisons between two independent groups, a two-tailed unpaired Student’s t-test was employed. For comparisons involving three or more groups, one-way analysis of variance (ANOVA) was conducted, followed by Dunnett’s post-hoc test to determine specific group differences. Statistical significance was denoted as follows: *p < 0.05, **p < 0.01, ***p < 0.001, and ****p < 0.0001. Results that did not achieve statistical significance were labeled as “ns” (not significant) or remained unmarked.

## 3 RESULTS

### 3.1 Determining interaction effects of aroma compounds by 3-AFC and S-curve

The phenomenon of olfactory interaction between 2,3,5-trimethylpyrazine (exhibiting a bakery-like odor) and L-menthol (characterized by an herbal aroma) was first demonstrated through designed sensory experiments. In this section, the olfactory detection thresholds were determined for both the individual compounds (2,3,5-trimethylpyrazine and L-menthol) and the binary mixture, enabling their interaction effects to be systematically compared (Fig. 1; Table 1). 2,3,5-trimethylpyrazine and L-menthol olfactory detection threshold concentrations were 10.75 mmol/L and 46.84 mmol/L, respectively (Fig. 1 a, d; Table S4). Subsequently, the thresholds of 2,3,5-trimethylpyrazine and L-menthol were examined again after equimolar mixing (each odorant maintains the same concentration as its single-substance solution). In the binary system, the detection thresholds were 81.32 mmol/L for 2,3,5-trimethylpyrazine and 275.49 mmol/L for L-menthol (Fig. 1 b, e; Table S4). Based on these results, equimolar mixing leads to a significant increase in the detection thresholds of both 2,3,5-trimethylpyrazine and L-menthol and their S-curves shifts to the right distinctly (Fig. 1 b, e). For 2,3,5-trimethylpyrazine, the measured detection threshold is larger than the theoretical detection threshold, with *D* (experimental threshold/theoretical threshold) =7.57, showing a masking effect (Table 1); for L-menthol, the measured detection threshold is larger than the theoretical detection threshold, with *D* (experimental threshold/theoretical threshold) =5.88, also showing a significant masking effect (Table 1).

**Fig. 1.**
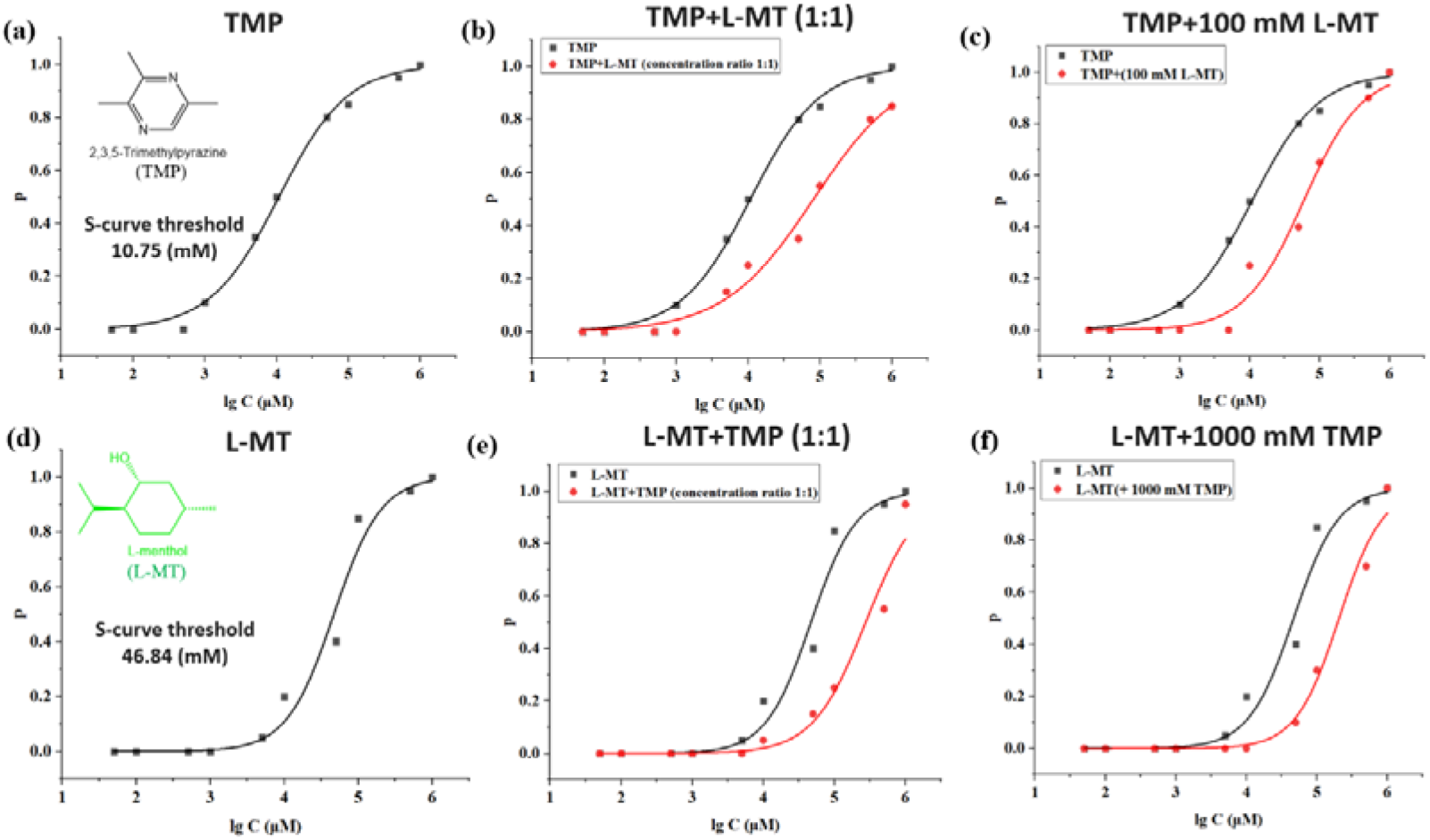
Results of olfactory threshold detection by the 3-AFC method. (a) and (d) represent the respective thresholds for 2,3,5-trimethylpyrazine and L-menthol, respectively; (c) represents the changes in 2,3,5-trimethylpyrazine after mixing with 100 mmol/L L-menthol; (b) and (e) represent the changes in the respective thresholds of 2,3,5-trimethylpyrazine and L-menthol after equimolar mixing; (f) represents the changes in L-menthol after mixing with 1000 mmol/L 2,3,5-trimethylpyrazine. TMP: 2,3,5-trimethylpyrazine, L-MT: L-menthol

**Table 1.**
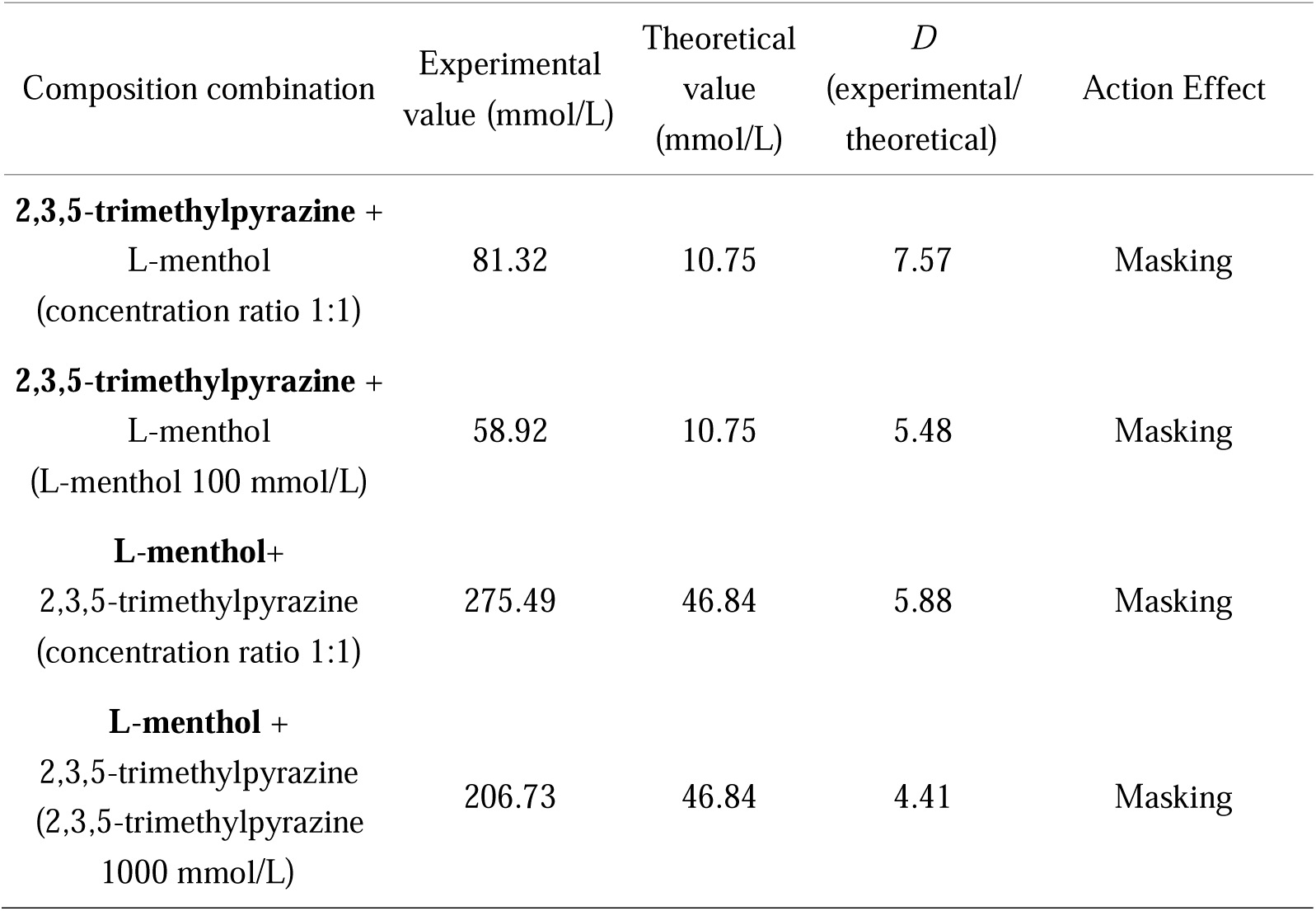
3-AFC-S curve discrimination of interactions between 2,3,5-trimethylpyrazine and L-menthol.

| Composition combination | Experimental<br>value (mmol/L) | Theoretical<br>value<br>(mmol/L) | *D*<br>(experimental/<br>theoretical) | Action Effect |
| --- | --- | --- | --- | --- |
| **2,3,5-trimethylpyrazine +**<br>L-menthol<br>(concentration ratio 1:1) | 81.32 | 10.75 | 7.57 | Masking |
| **2,3,5-trimethylpyrazine +**<br>L-menthol<br>(L-menthol 100 mmol/L) | 58.92 | 10.75 | 5.48 | Masking |
| **L-menthol+**<br>2,3,5-trimethylpyrazine<br>(concentration ratio 1:1) | 275.49 | 46.84 | 5.88 | Masking |
| **L-menthol +**<br>2,3,5-trimethylpyrazine<br>(2,3,5-trimethylpyrazine<br>1000 mmol/L) | 206.73 | 46.84 | 4.41 | Masking |

Initial mutual masking effects were characterized by the measurements of threshold in equimolar binary mixtures. To further investigate the interaction pattern, an alternative experimental approach was employed wherein the concentration of one odorant remained fixed in the binary mixture system while the detection threshold of another one was assessed. First, the concentration of L-menthol was fixed at 100 mmol/L, and the concentration of 2,3,5-trimethylpyrazine was varied to achieve L-menthol : 2,3,5-trimethylpyrazine =1000:1/200:1/100:1/20:1/10:1/2:1/1:1/1:5/1:10. The olfactory detection thresholds of 2,3,5-trimethylpyrazine was measured as 58.92 mmol/L and the measured detection threshold was larger than the theoretical detection threshold, with *D* (experimental threshold/theoretical threshold) =5.48, showing a masking effect. In turn, to investigate the effect of 2,3,5-trimethylpyrazine on L-menthol, the 2,3,5-trimethylpyrazine concentrations were fixed at 1000 mmol/L, and the concentration ratio of 2,3,5-trimethylpyrazine to L-menthol was varied to achieve 2,3,5-trimethylpyrazine : L-menthol=10000:1/2000:1/1000:1/200:1/100:1/20:1/10:1/2:1/1:1. The thresholds of L-menthol were measured to be 206.73 mmol/L and the measured detection threshold was larger than the theoretical detection threshold, *D* (experimental threshold/theoretical threshold) =4.41, which also showed a masking effect (Fig. 1 c, f; Table 1).

### 3.2 Effect of odorants on cell viability

Research on odorant interactions based on olfactory receptors should be conducted using appropriate concentrations of odorants. Therefore, 24 hours post-transfection, odorants at varying concentrations were applied to cells for viability assessment. In this experiment, the odorant treatment duration was set at 10 minutes, which aligns with the time frame for cAMP signal detection following ligand application. The results demonstrated that 2,3,5-trimethylpyrazine exhibited no cytotoxicity within the 0-1000 μmol/L range in OR5K1-expressing cells (Fig. 2 a). However, in groups containing L-menthol - whether administered alone or in 1:1 combination with 2,3,5-trimethylpyrazine - concentrations exceeding 100 μmol/L showed significant cellular toxicity (Fig. 2 b, c). OR2W1-expressing cells displayed an identical response pattern (Fig. 2 d-f). Consequently, subsequent experiments employed concentration ranges of 1-1000 μmol/L for 2,3,5-trimethylpyrazine and 1-100 μmol/L for L-menthol administration.

**Fig. 2.**
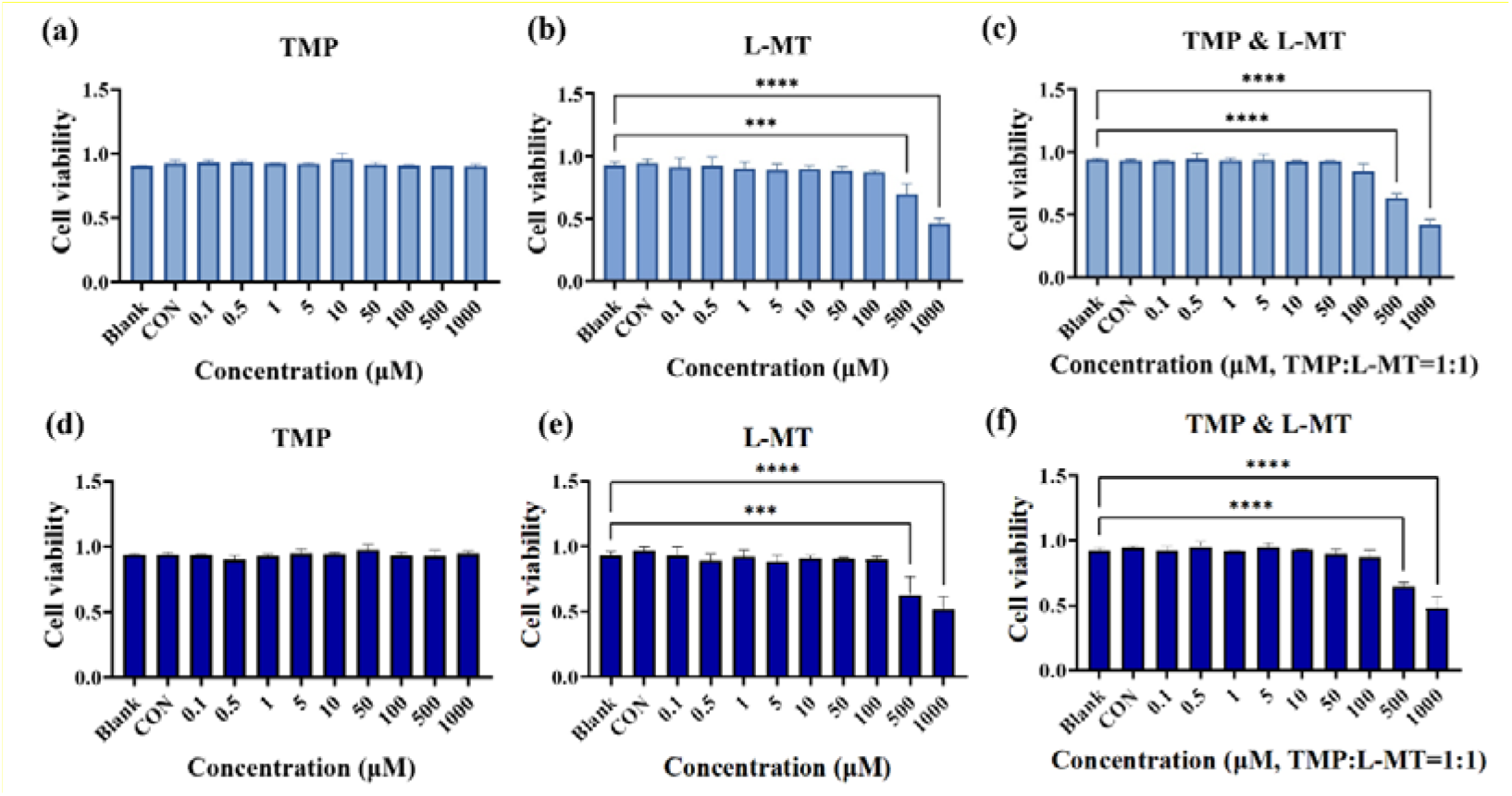
Determination of cell viability after application of odorants. (a)-(c) represent the cell viability of OR5K1-expressing cells treated with varying concentrations of 2,3,5-trimethylpyrazine, L-menthol, and 2,3,5-trimethylpyrazine+L-menthol, respectively; (d)-(f) represent the cell viability of OR2W1-expressing cells treated with varying concentrations of 2,3,5-trimethylpyrazine, L-menthol, and 2,3,5-trimethylpyrazine+L-menthol, respectively. The “Blank” group represents no treatment was performed; the “CON” group represents the treatment with DMSO solvent alone. Statistical significance was denoted as follows: *p < 0.05, **p < 0.01, ***p < 0.001, and ****p < 0.0001. Results that did not achieve statistical significance were labeled as “ns” (not significant) or remained unmarked. TMP: 2,3,5-trimethylpyrazine, L-MT: L-menthol

### 3.3 OR5K1 is a responsive receptor for 2,3,5-trimethylpyrazine

A modified *in vitro* olfactory receptor screening assay was employed to evaluate human olfactory responses to 2,3,5-trimethylpyrazine in this study. Systematic screening of a panel of 294 ORs (excluding mutant variants), followed by quantitative analysis of their response profiles, was conducted at two distinct concentrations of 2,3,5-trimethylpyrazine (100 and 1000 μmol/L). The results demonstrate that OR5K1 is a responsive receptor for 2,3,5-trimethylpyrazine, regardless of whether the concentration is 100 μmol/L or 1000 μmol/L (Fig. 3 a, b). Accordingly, OR5K1 serves as an ideal target for studying 2,3,5-trimethylpyrazine-other odorants interactions and its robust response to 2,3,5-trimethylpyrazine enables systematic investigation of odor mixture modulation mechanisms at the receptor level.

**Fig. 3.**
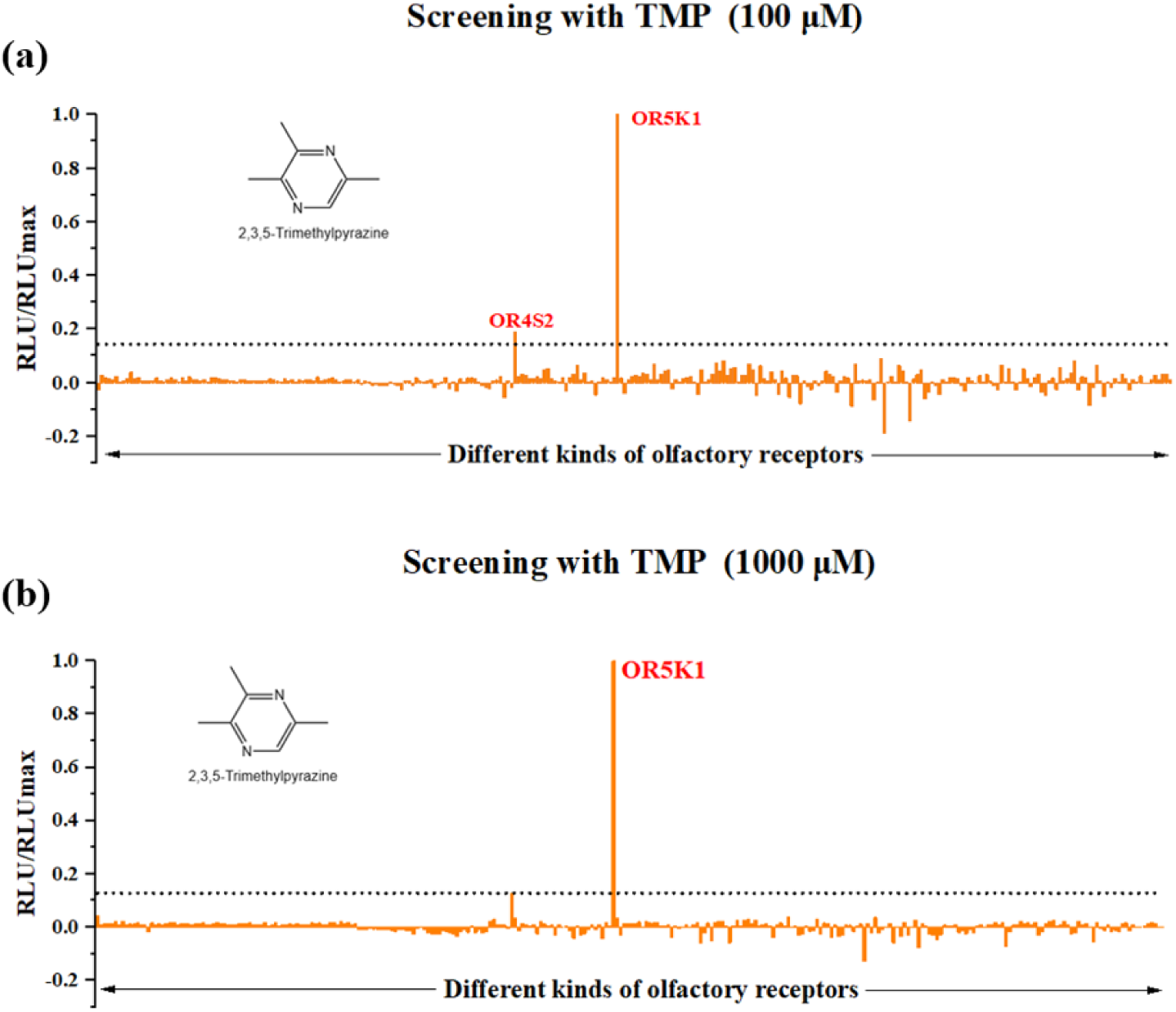
2,3,5-trimethylpyrazine exclusively activates OR5K1. 294 human ORs were screened with 100 μmol/L 2,3,5-trimethylpyrazine (a) and 1000 μmol/L 2,3,5-trimethylpyrazine (b). The response values for each receptor are shown (n=1 measured in duplicate) and the dashed line indicates the 2σ threshold (2-fold the standard deviation above the average of all signaling amplitudes in a screening experiment). Data were mock control-subtracted and normalized to the maximum response. RLU = relative luminescence unit. TMP: 2,3,5-trimethylpyrazine

The concentration-response curve serves as a fundamental analytical approach in olfactory research, providing critical information for both validating receptor-ligand interactions and determining half-maximal effective concentrations (EC_50_) of odorants ^41^. Therefore, the concentration-response relationship of 2,3,5-trimethylpyrazine and L-menthol on OR5K1 was constructed and quantitative analysis revealed a dose-dependent activation pattern of OR5K1 by 2,3,5-trimethylpyrazine, with response amplitudes progressively increasing across a concentration range of 0.1-1000 μmol/L. Nonlinear regression analysis yielded an EC_50_ value of 27.67±2.02 μmol/L for 2,3,5-trimethylpyrazine (Fig. 4 a). In contrast, L-menthol didn’t elicit significant receptor activation at any tested concentration, as evidenced by the absence of detectable response signals and the inability to establish meaningful concentration-response relationships (Fig. 4 a).

**Fig. 4.**
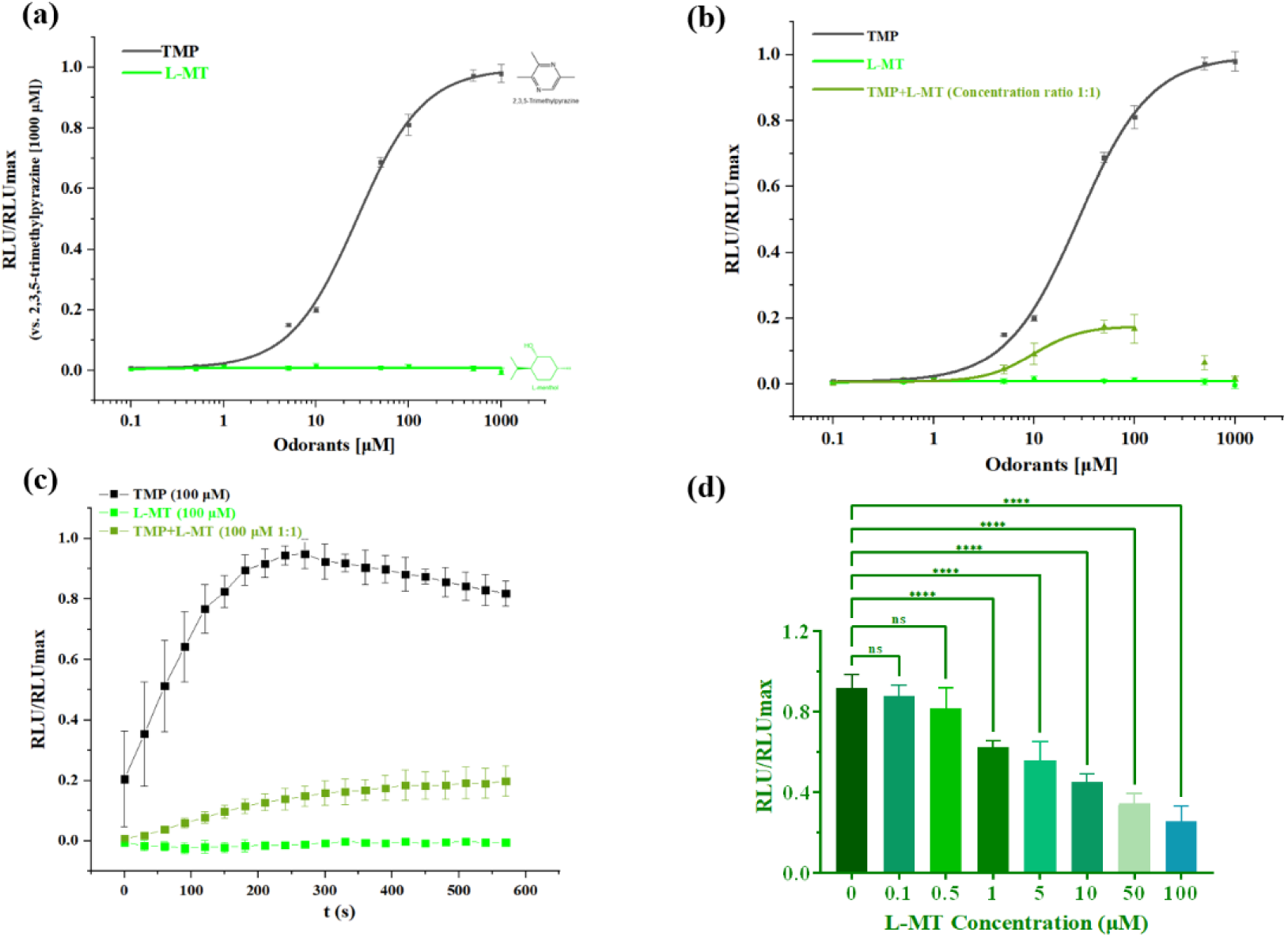
Concentration-response curves for individual and binary systems on OR5K1. (a) represents the concentration-response curves for 2,3,5-trimethylpyrazine and L-menthol on OR5K1; (b) represents the concentration-response curves for 2,3,5-trimethylpyrazine, L-menthol, and 2,3,5-trimethylpyrazine+L-menthol (in equimolar concentrations); (c) represents the time-response curves for 2,3,5-trimethylpyrazine, L-menthol, and 2,3,5-trimethylpyrazine+L-menthol (in equimolar concentrations) at 100 μmol/L; (d) represents fixing the 2,3,5-trimethylpyrazine concentration and varying the concentration of L-menthol. All data were mock control-subtracted and normalized to the maximum response, and are shown as mean ± SD (n = 4). RLU = relative luminescence unit. Statistical significance was denoted as follows: *p < 0.05, **p < 0.01, ***p < 0.001, and ****p < 0.0001. Results that did not achieve statistical significance were labeled as “ns” (not significant) or remained unmarked. TMP: 2,3,5-trimethylpyrazine, L-MT: L-menthol

### 3.4 Binary interactions between L-menthol and 2,3,5-trimethylpyrazine based on OR5K1

Previous psychophysical studies have demonstrated that eatable alkanols specifically L-menthol could modulate the olfactory perception of 2,3,5-trimethylpyrazine (Fig. 1; Table 1). However, the underlying mechanisms remain unexplored at the cellular level. In the present study, mechanistic evidence is provided for how L-menthol influences 2,3,5-trimethylpyrazine detection at the single-receptor level, elucidating its modulatory effects on receptor activation dynamics.

In our experiments, L-menthol was mixed with 2,3,5-trimethylpyrazine at the same concentration ratio to obtain a series of binary systems of L-menthol and 2,3,5-trimethylpyrazine with different concentration gradients, such as 0.1 μmol/L L-menthol+0.1 μmol/L 2,3,5-trimethylpyrazine, 0.5 μmol/L L-menthol+0.5 μmol/L 2,3,5-trimethylpyrazine, and so on (see Fig. S2 for details). The results obtained from GC-Q/Orbitrap MS analysis of the odorants and their mixed systems demonstrated the absence of any new compound peaks, suggesting that no chemical reaction occurred between the binary odorants to generate novel substances. This ensures that the mutual inhibitory effect between the compounds arises from a physiological response rather than a chemical interaction of the mixtures (Fig. S3). Compared with the 2,3,5-trimethylpyrazine treatment group, the response level of cells in equimolar concentrations of L-menthol and 2,3,5-trimethylpyrazine was significantly reduced, and this inhibition was present at each concentration (Fig. 4 b). On the time-response curve, the response values of the binary system (100 μmol/L 2,3,5-trimethylpyrazine+100 μmol/L L-menthol) were more than 80% lower than those of the 2,3,5-trimethylpyrazine-treated group (Fig. 4 c).

In fact, only making L-menthol mixed with 2,3,5-trimethylpyrazine in equimolar concentrations was not enough to obtain convincing conclusions, so the concentration ratios between 2,3,5-trimethylpyrazine and L-menthol were varied to observe their effects on the process of OR5K1 activation. Compared with the 1000 μmol/L 2,3,5-trimethylpyrazine-treated group, the presence of L-menthol inhibited the OR5K1 response signals remarkably. The interaction of L-menthol on 2,3,5-trimethylpyrazine-OR5K1 was concentration-dependent within 0.1-100 μmol/L. When L-menthol concentrations are more than 1 μmol/L, the signals began to decrease significantly compared to the 1000 μmol/L 2,3,5-trimethylpyrazine-treated group, and this inhibitory effect was even more pronounced with increasing L-menthol concentrations (Fig. 4 d). This concentration-inhibition relationship provides experimental validation for L-menthol’s suppressive effect on 2,3,5-trimethylpyrazine, effectively excluding random or artifactual influences. The concentration-dependent weakening on receptor activation induced by 2,3,5-trimethylpyrazine confirms the physiological inhibition of L-menthol.

### 3.5 Binary interactions between 2,3,5-trimethylpyrazine and L-menthol based on OR2W1

As OR2W1 responds broadly to key food odorants (aldehydes, alcohols, ketones) and to L-menthol (not 2,3,5-trimethylpyrazine), it was selected as an additional key receptor to verify 2,3,5-trimethylpyrazine’s influence on L-menthol-induced agonism. Our experimental results showed that the response of OR2W1 to L-menthol was concentration-dependent from 0.1-100 μmol/L, with an EC_50_ value of 22.69±3.85 μmol/L (Fig. 5 a).

**Fig. 5.**
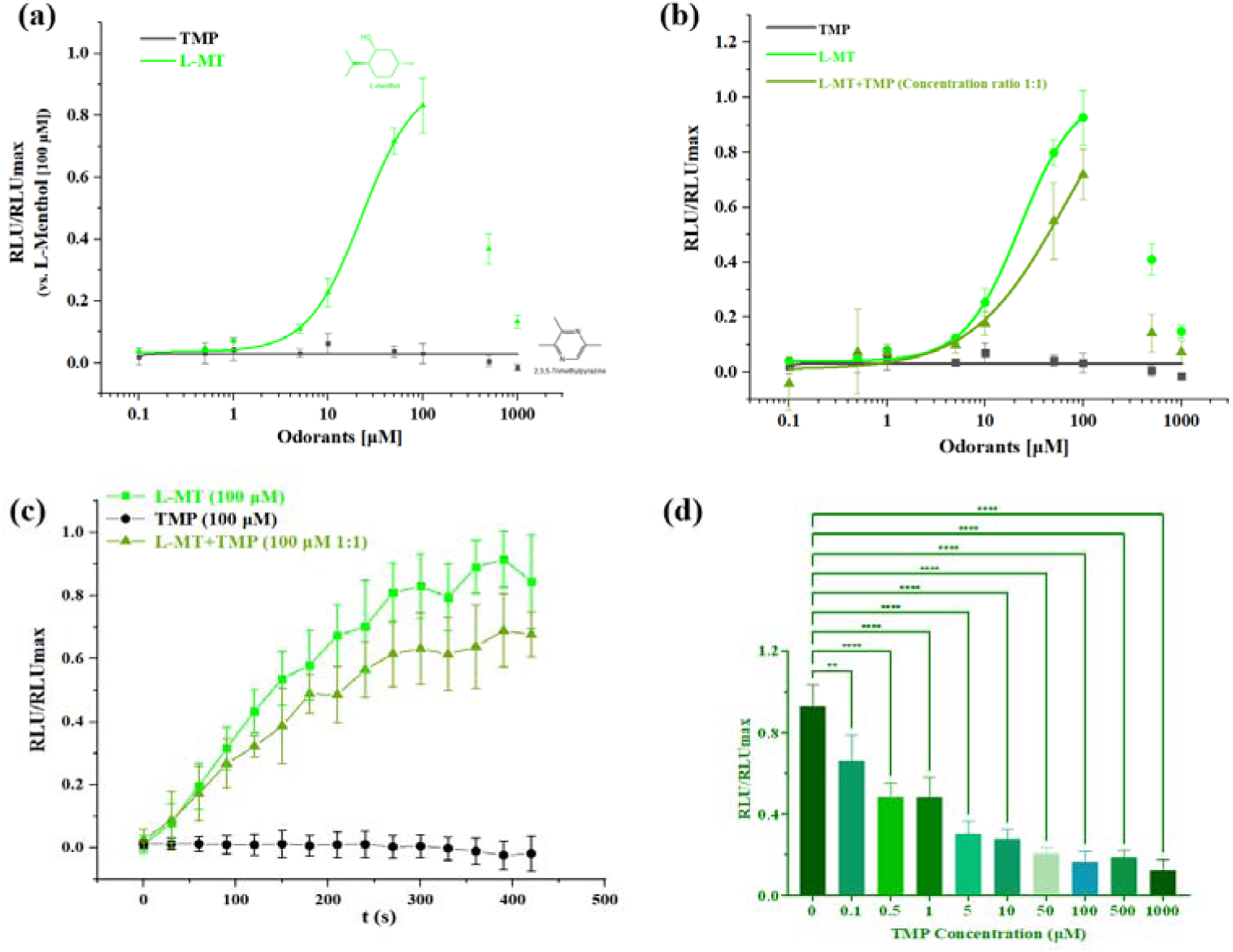
Concentration-response curves for individual and binary systems on OR2W1. (a) concentration-response curves for L-menthol and 2,3,5-trimethylpyrazine on OR2W1; (b) represents the concentration-response curves for L-menthol, 2,3,5-trimethylpyrazine, and L-menthol+2,3,5-trimethylpyrazine (in equimolar concentrations); (c) represents the time-response curves for L-menthol, 2,3,5-trimethylpyrazine, L-menthol+2,3,5-trimethylpyrazine (in equimolar concentrations); (d) represents fixing the L-menthol concentration and varying the concentration of 2,3,5-trimethylpyrazine. All data were mock control-subtracted and normalized to the maximum response, and are shown as mean ± SD (n = 4). RLU = relative luminescence unit. Statistical significance was denoted as follows: *p < 0.05, **p < 0.01, ***p < 0.001, and ****p < 0.0001. Results that did not achieve statistical significance were labeled as “ns” (not significant) or remained unmarked. TMP: 2,3,5-trimethylpyrazine, L-MT: L-menthol

The ligand that OR2W1 respond to is L-menthol but not 2,3,5-trimethylpyrazine, so the core substances investigated on this receptor was L-menthol. In the presence of the equimolar concentrations of 2,3,5-trimethylpyrazine, the activation of OR2W1 by L-menthol was reduced (Fig. 5 b, c) and there was a concentration dependence of the response of the binary system in the range of 0.1-100 μmol/L that is similar to the results demonstrated on OR5K1. To investigate the concentration-dependent modulation by 2,3,5-trimethylpyrazine, L-menthol was maintained at a constant concentration of 100 μmol/L while 2,3,5-trimethylpyrazine levels were varied from 0.1 to 1000 μmol/L to assess the modulatory effects on OR2W1 activation dynamics. The results showed that 2,3,5-trimethylpyrazine had a concentration-dependent physiological inhibitory effect on the activation of OR2W1 by L-menthol (Fig. 5 d). This suggests that the two odorants will inhibit each other from the receptor perspective of 2,3,5-trimethylpyrazine and L-menthol.

### 3.6 Molecular modeling predicts binary odorant interactions at the OR level

Assays using olfactory receptor-expressing cell systems have validated the functional interaction between 2,3,5-trimethylpyrazine and L-menthol. However, the precise molecular mechanisms remain unclear, prompting the application of computational molecular modeling to investigate the potential modulation mechanisms of diverse odorants on olfactory receptor activation. AlphaFold3 was used to predict the co-folding of small molecules and proteins, which could directly predict binding sites and modes (Fig. 6 a, b). employed in our recently published study and are included here for comparison ^42^. For OR2W1, molecular docking simulations revealed that the hydroxyl groups of L-menthol could form stable hydrogen bonds with the Y259^6.55^. 2,3,5-trimethylpyrazine forms only weak hydrophobic interactions with Y259^6.55^ in OR2W1, insufficient to trigger activation (Fig. 6 c, d). However, for OR5K1, 2,3,5-trimethylpyrazine establishes hydrophobic interaction with L255^6.51^, hydrogen bonding with S203^5.43^ and π-π stacking with H155^4.56^, potentially constituting its activation mechanism, while L-menthol lack these comprehensive interactions, preventing receptor activation (Fig. 6 e, f).

**Fig. 6.**
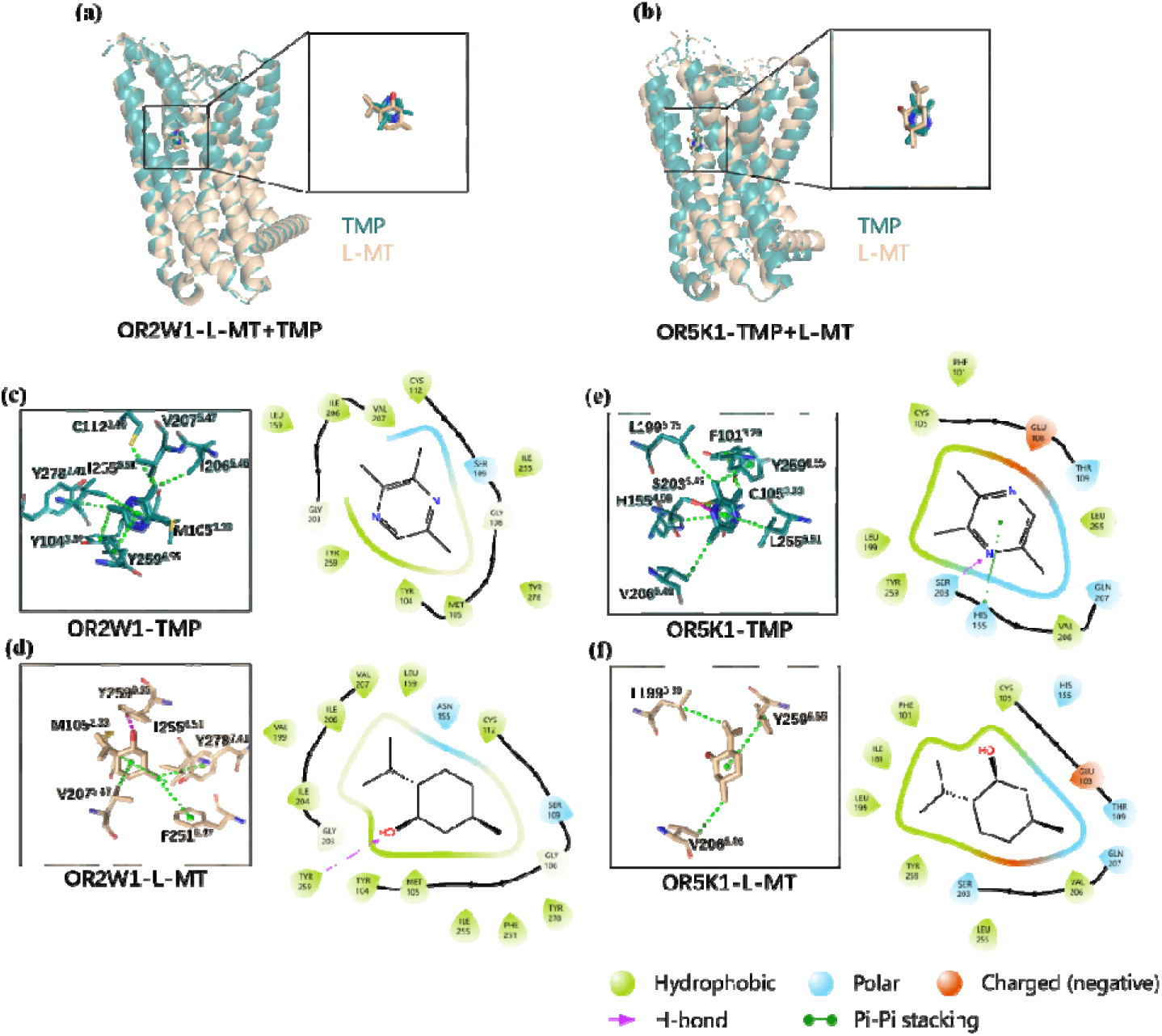
Molecular docking shows interactions between odorants and olfactory receptors. (a) represents the overlapping state in which L-menthol and 2,3,5-trimethylpyrazine co-interact with OR2W1; and (b) represents the overlapping state in which 2,3,5-trimethylpyrazine and L-menthol co-interact with OR5K1; (c) and (d) represent OR2W1 interaction modes with 2,3,5-trimethylpyrazine and L-menthol, respectively; (e) and (f) represent OR5K1 interaction modes with 2,3,5-trimethylpyrazine and L-menthol, respectively. In the 3D diagram, the green dashed line indicates hydrophobic interactions and the purple dashed line indicates hydrogen bonding. TMP: 2,3,5-trimethylpyrazine, L-MT: L-menthol

Based on the AlphaFold 3-predicted structures, 200 ns molecular dynamics (MD) simulations were performed for all odorant-receptor systems ^42^. First, the MM-PBSA method was employed to calculate binding free energies (Table 2). The results showed that OR2W1 exhibited lower binding energy with L-menthol than with 2,3,5-trimethylpyrazine, suggesting higher affinity for L-menthol. In contrast, 2,3,5-trimethylpyrazine showed the lowest binding energy and highest affinity for OR5K1. Further decomposition of binding free energies identified the top 10 contributing residues. The key residues of OR2W1 (e.g., M105^3.33^, I255^6.51^) and OR5K1 (S203^5.43^, V206^5.46^, L255^6.51^, L199^5.39^) showed interactions consistent with AlphaFold 3 predictions (Fig. 6; Fig. 7 a, b). Additionally, the root-mean-square deviation (RMSD) of ligand heavy atoms was calculated following alignment of protein Cα atoms using the AlphaFold 3-predicted structure as a reference (Fig. 7 c, d). For OR2W1, L-menthol maintained an RMSD of ∼0.5 nm, indicating stable binding, while 2,3,5-trimethylpyrazine displayed larger RMSD fluctuations and eventually dissociated from the binding pocket. In OR5K1, both ligands remained bound initially, but L-menthol departed from its original binding site around 200 ns, whereas 2,3,5-trimethylpyrazine binding remained relatively stable (Fig. 7 c, d). In both receptor systems, 2,3,5-trimethylpyrazine and L-menthol exhibit similar interaction patterns, suggesting competitive binding at the active pocket. Their mutual antagonism likely results from analogous binding modes preventing co-occupancy. These computational findings corroborate both cellular assays and molecular modeling results, collectively showing the reliability of AlphaFold 3 co-folding predictions.

**Fig. 7.**
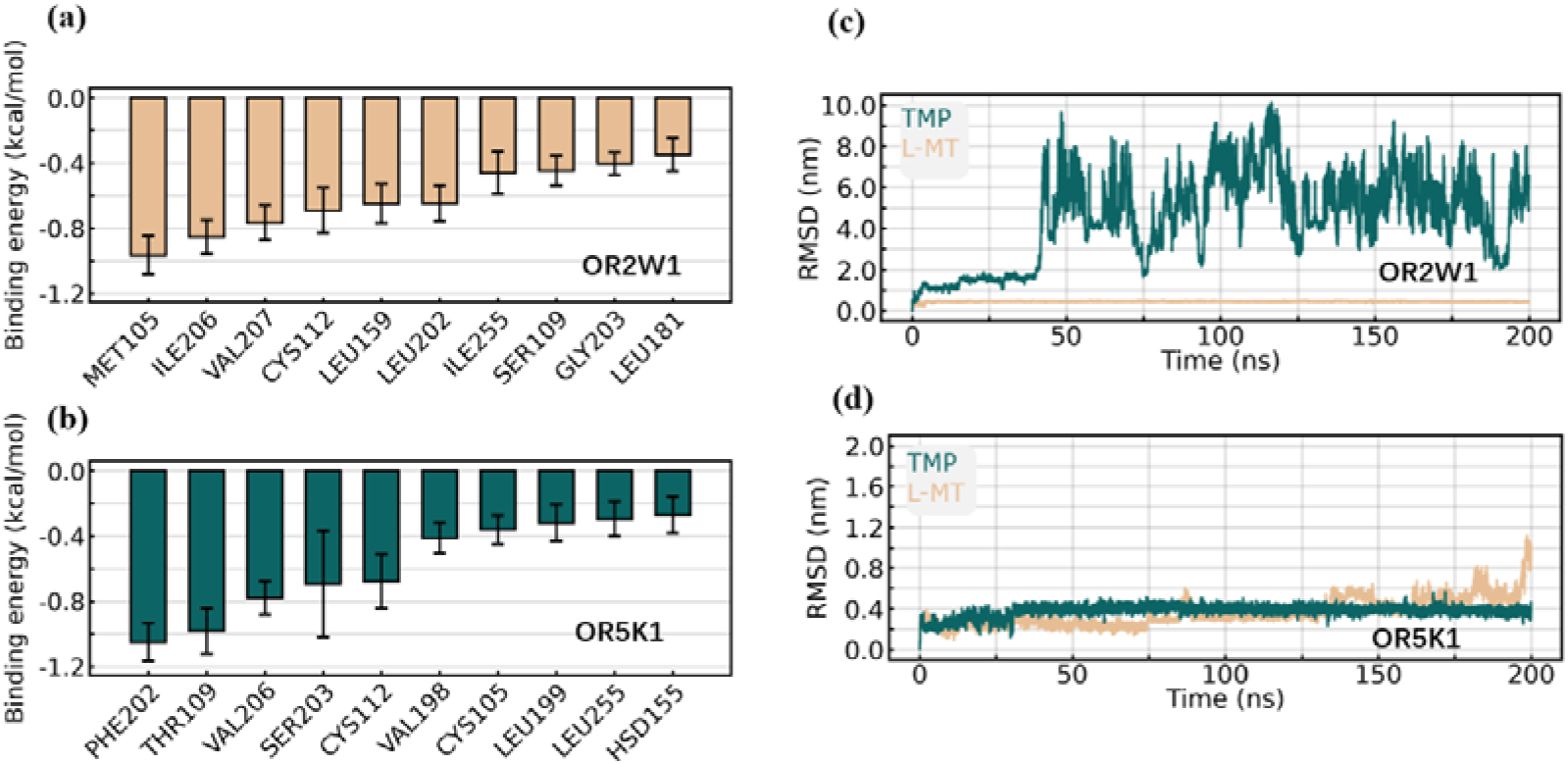
Molecular dynamics shows binding of odorants and olfactory receptors. (a) represents the 10 most important key residues in the interaction of L-menthol with OR2W1; (b) represents the 10 most important key residues in the interaction of 2,3,5-trimethylpyrazine with OR5K1; (c) represents the RMSD of 2,3,5-trimethylpyrazine (green), L-menthol (yellow), and OR2W1 within 200 ns; and (d) represents the RMSD of 2,3,5-trimethylpyrazine (green), L-menthol (yellow), and OR5K1 within 200 ns. TMP: 2,3,5-trimethylpyrazine, L-MT: L-menthol

**Table 2.** Docking binding energy of olfactory receptors to odorants.

| $\Delta G$ Mean ± SEM<br>(kcal/mol) | L-menthol | 2,3,5-trimethylpyrazine |
| --- | --- | --- |
| OR2W1 | **-17.21 ± 0.08** | -2.65 ± 0.09 |
| OR5K1 | -4.80 ± 0.40 | **-13.34 ± 0.09** |

## 4 DISCUSSION

In the sensory evaluation of food flavors, common methods include the duo-trio test, two-out-of-five test, and 3-AFC test. Among these, the 3-AFC test combined with the S-curve model demonstrates outstanding performance in terms of stability and accuracy and has been applied to flavor analysis across various food categories ^43–45^. Through this method, our results demonstrate suppressed olfactory perception of 2,3,5-trimethylpyrazine in the presence of L-menthol, explaining the diminished baked aromas observed with menthol-containing spices. Furthermore, bidirectional masking between 2,3,5-trimethylpyrazine and L-menthol was observed (Fig. 1 and Table 1), consistent with the pattern reported for many binary aromatic compounds ^28^. However, sensory evaluation based on subjective analysis is inherently limited, which may induce variability in sensory data. Such limitations involve the chemical complexity of odorants, the complexity of human perception, the susceptibility of human panelists to fatigue during long-term tests and psychological biases in judgment ^46^. In addition, odor samples for sensory experiments are prepared via solution mixing in the majority of published studies on odorants interaction ^43–46^. Despite the simplicity of this method, the volatility of mixed solutions containing different odorants may undergo alterations. In other words, the combination of different odorants may cause changes in the concentration of individual components, with more significant variations observed in the concentration of gaseous components after volatilization. Therefore, direct gaseous mixing of odorants is the most suitable candidate method. Nevertheless, the implementation of dynamic gaseous mixing and evaluation with precise concentrations is challenging due to the constraints of experimental conditions. For this reason, our experiments achieve the mixing of gaseous substances through the volatilization of their solutions in a non-contact system (separate filter papers), thus avoiding the uncertainty caused by the direct mixing of solutions. Odorant concentration measurements using a headspace-based GC-MS method showed that the odor concentrations resulting from the mixed addition of filter papers containing different odorants were nearly identical to those from the addition of filter papers with a single odorant alone. This suggests that the mixing effect of gaseous odorants does not affect the volatility of individual odorants (Fig. S4). Despite such precautions, certain inherent uncertainties persist, including the inability to standardize the volatilization efficiency of odorants from filter papers and the interference introduced by the opening and closing of vial lids during experimental operation. Subsequent research will explore more optimized and simplified approaches for odor sample preparation to refine olfactory assessment protocols, encompassing the development of direct gas-phase premixing technologies and the establishment of an integrated system that couples odor preparation with quantitative headspace analysis ^47, 48^.

Cellular assays using OR-expressing systems validated sensory findings. The first step involved screening for olfactory receptors responsive to 2,3,5-trimethylpyrazine. Interestingly, in addition to the identified OR5K1, the results revealed that OR4S2 was also responsive at 100 μmol/L 2,3,5-trimethylpyrazine, highlighting the differential response patterns of ORs across varying concentrations (Fig. 3 a). However, despite this, the maximum response value of OR4S2 is still far from OR5K1, so it was not considered in this study (Fig. 3 a). OR5K1, established as a broad-spectrum pyrazine detector ^49^, was selected to study L-menthol’s effect on 2,3,5-trimethylpyrazine response. OR2W1, responsive to L-menthol, was used to investigate 2,3,5-trimethylpyrazine’s effect on L-menthol activity. A bidirectional mutual inhibitory effect between 2,3,5-trimethylpyrazine and L-menthol was presented at both receptors (Fig. 4 b; Fig. 5 b). Notably, 2,3,5-trimethylpyrazine exerted weaker suppression on L-menthol-mediated OR2W1 activation than L-menthol did on 2,3,5-trimethylpyrazine-mediated OR5K1 activation. This difference may stem from OR2W1’s functional characteristics. Although OR2W1 is not activated by 2,3,5-trimethylpyrazine, its broad tuning suggests that its binding pocket is potentially promiscuous, favoring compatibility with different odorants ^33, 50–52^. This contrasts with OR5K1’s high selectivity for 2,3,5-trimethylpyrazine, indicating a more constrained binding architecture ^53^. Consequently, L-menthol may induce greater perturbation in the precise OR5K1-2,3,5-trimethylpyrazine interaction. Additionally, the exploration of the relationships between olfactory receptors and odorants constitutes a crucial approach for investigating the odor perception mechanisms and survival rules of humans and animals in nature. With respect to olfactory receptors OR5K1 and OR2W1, previous studies have been conducted by other researchers: for instance, Marcinek *et al.* demonstrated that OR5K1 is a conserved olfactory receptor dedicated to the discrimination of pyrazine derivatives across mammalian species ^49^, and Haag *et al.* identified that OR2W1 functions as a broadly tuned receptor responding to key aroma compounds in Western foods ^33^. These studies on olfactory receptors based on physiological research models are pioneering in the field. However, the aroma components in foods form a complex and extensive system, and human perception of odorants is largely dependent on the interactions among odorant molecules. In the present study, we focused on two common aroma compounds in thermally processed and seasoned foods, namely 2,3,5-trimethylpyrazine and L-menthol, and adopted heterologous expression cell models of their corresponding responsive olfactory receptors (OR5K1 and OR2W1) to investigate the interaction mode between the two compounds. This study can be regarded as an extension and practical application of cell biological experiments, and also provides a research paradigm for investigating the interaction effects of aroma compounds based on olfactory receptors.

Visualizing the results of physiological experiments via molecular modeling is one of the universal approaches for inferring the underlying mechanisms of action. In human ORs, hydrogen bonding between the compound hydroxyl group and the residues of the receptor could change the shape of the active pocket, which in turn alters the conformation of the transmembrane sequence and extracellular rings ^54, 55^. AlphaFold3 predictions suggest 2,3,5-trimethylpyrazine and L-menthol competitively bind to key pockets in OR5K1 and OR2W1, respectively, with mutual inhibition likely resulting from competition at critical residues (Fig. 6). Specifically, in OR2W1, 2,3,5-trimethylpyrazine forms hydrophobic interactions with Y259^6.55^ (a residue previously shown to preferentially interact with nonpolar molecules ^56, 57^) which may disrupt the hydrogen bonding network required for L-menthol-mediated activation (Fig. 6 c, d). Similarly, while 2,3,5-trimethylpyrazine activates OR5K1 through interactions with critical residues (L255^6.51^, S203^5.43^, L199^5.39^), L-menthol appears to sterically hinder this activation process when binding to these same residues (Fig. 6 e, f; Fig. 7 a, b). Molecular dynamics simulations further revealed that 2,3,5-trimethylpyrazine exhibits relatively unstable binding in OR2W1’s pocket compared to L-menthol’s more stable occupation of OR5K1’s binding site (Fig. 7 c, d). The stable occupation of non-activating ligands within orthosteric binding pockets can potentially interfere with agonist-induced receptor activation ^53^. This mechanistic insight explains why L-menthol exhibits stronger inhibitory effects on 2,3,5-trimethylpyrazine-mediated OR5K1 activation compared to 2,3,5-trimethylpyrazine’s inhibition of L-menthol-OR2W1 signaling. Furthermore, molecular simulation studies have revealed that compared to individual odorants, mixed odorant systems often exhibit higher binding energies and less stable binding conformations ^58^. It provides additional mechanistic insights into inter-odorant interactions. These findings collectively establish a structure-activity relationship underlying the reciprocal inhibition between these two odorants at their respective target receptors. While computational approaches are widely employed, current computer-predicted models of olfactory receptor activation remain incomplete. Comprehensive understanding requires experimental validation through site-directed mutagenesis and cryo-EM structural determination, particularly for characterizing odorant interactions regarding their competitive/non-competitive binding modes and conformational effects on receptor domains ^54^.

In conclusion, masking effect between aroma substances is widespread in natural food systems and food processing. Our study systematically elucidates the mutual masking effects between thermally generated 2,3,5-trimethylpyrazine and spice-derived L-menthol through a multi-level approach combining sensory evaluation, cellular assays, and molecular simulations. To scientifically determine the human perception of the interaction between 2,3,5-trimethylpyrazine and L-menthol, a sensory evaluation was conducted using the 3-AFC approach. The results preliminarily confirm that TMP and L-menthol exhibit mutual masking effects. And then, receptor screening revealed OR5K1 as a highly responsive receptor for 2,3,5-trimethylpyrazine, where L-menthol exhibited significant suppression of 2,3,5-trimethylpyrazine response signal. Conversely, results demonstrated an inhibitory effect of 2,3,5-trimethylpyrazine on L-menthol perception via OR2W1, with cellular-level findings consistent with sensory evaluation results. Molecular docking and dynamics simulations provided structural insights, showing that both odorants share similar binding modes in their respective receptors, suggesting competitive binding as the underlying mechanism. This comprehensive investigation offers profound mechanistic understanding of flavor interactions in food systems, with substantial practical implications for food flavor design and optimization. Nevertheless, the current research still has certain limitations. ORs represent primary detectors, while holistic olfactory perception involves multidimensional complexity. Discrepancies between receptor signaling and integrated perception arise from combinatorial coding in human olfaction ^1^. Therefore, sensory and cellular experiments are complementary. A comprehensive understanding of 2,3,5-trimethylpyrazine interactions requires: (1) systematic analysis of all responsive receptors, and (2) translational validation at the organismal level (e.g., primary olfactory tissue and murine models). This could involve measuring electrophysiological responses and biochemical changes in isolated neurons or relevant brain regions using techniques like high-speed 3D imaging or optogenetics ^59, 60^. Furthermore, while heterologous expression coupled with second messenger (e.g., cAMP) detection enables high-throughput OR screening, challenges include potential signaling pathway ambiguity and interference from endogenous host cell receptors ^61^. Deeper biological investigations are needed.

## Abbreviations

TMP: 2,3,5-trimethylpyrazine
L-MT: L-menthol
GC-O-MS: gas chromatography-olfactometry-mass spectrometry
ORs: olfactory receptors
FBS: fetal bovine serum
DiPG: dipropylene glycol
3-AFC: three-alternative forced-choice
PCR: polymerase chain reaction
DMSO: dimethyl sulfoxide

## Acknowledgments

This work was supported by grants from the Key Program of the National Natural Science Foundation of China Regional Innovation and Development Joint Fund (Grant No. U24A20474), the Key Program of National Natural Science Foundation of China (Grant No. 32330080), and Shanghai Jiaotong University new faculty initiation program (Grant No. 24X010502905).

## Supporting Information

Information for *Homo sapiens* OR5K1 and OR2W1 (Table S1); information for all ORs used for screening (excluding mutant variants) (Table S2); interpretation of units of measure (Table S3); calculate the threshold value based on the S-curve method (Table S4); information for the vector of target gene plasmids (Figure S1); a cellular experimental design protocol for exploring binary interactions of odorants (Figure S2); composition testing of binary systems (Figure S3); the concentration of odorants in the gaseous mixture system determined by GC-MS method (Figure S4); and detailed instructions for sensory experiments.

## TOC

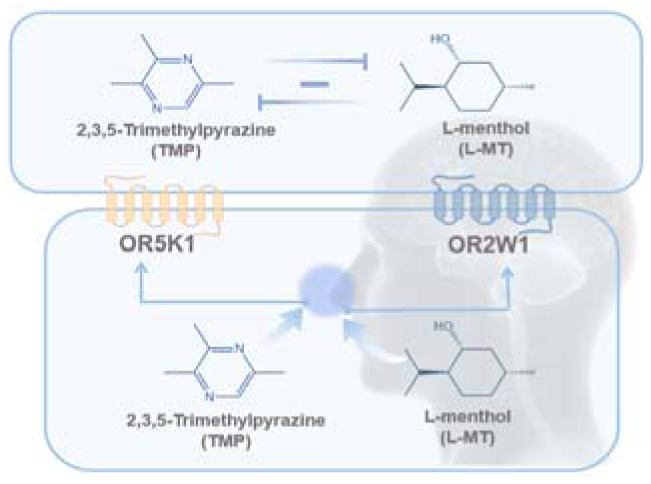

## Notes

### Competing Interest Statement

The authors have declared no competing interest.

### Summary of Updates

Our manuscript has been accepted; we will update it with the latest version first.

