## Supporting Information for "Elucidating Odorant Masking Effects in Food Matrices Through Olfactory Receptor-Based Profiling" (from another version of this preprint: version 1, 2026-01-15)

| **Gene Name** | **Rho-OR5K1** |
| --- | --- |
| **Gene Sequence** | ACGCGTATGAACGGCACCGAGGGCCCAAACTTCTACGTGCCTTTCTCCAACAAGACAGGAGTGGTCATGGCTGAAGAAAATCATACCATGAAAAATGAGTTTATCCTCACAGGATTTACAGATCACCCTGAGCTGAAGACTCTGCTGTTTGTGGTGTTCTTTGCCATCTATCTGATCACCGTGGTGGGGAATATTAGTTTGGTGGCACTGATATTTACACACCGTCGGCTTCACACACCAATGTACATCTTTCTGGGAAATCTGGCTCTTGTGGATTCTTGCTGTGCCTGTGCTATTACCCCCAAAATGTTAGAGAACTTCTTTTCTGAGAACAAAAGGATTTCCCTCTATGAATGTGCAGTACAGTTTTATTTTCTTTGCACTGTGGAAACTGCAGACTGCTTTCTTCTGGCAGCAATGGCCTATGACCGCTATGTGGCCATATGCAACCCACTGCAGTACCACATCATGATGTCCAAGAAACTCTGCATTCAGATGACCACAGGGGCCTTCATAGCTGGAAACCTGCATTCCATGATTCATGTAGGGCTTGTATTTAGGTTAGTTTTCTGTGGATCGAATCACATCAACCACTTTTACTGTGATATTCTTCCCTTGTATAGACTCTCTTGTGTTGATCCTTATATCAATGAACTGGTTCTATTCATCTTCTCAGGTTCAGTTCAAGTCTTTACCATAGGTAGTGTCTTAATATCTTATCTCTATATTCTTCTTACTATTTTCAAAATGAAATCCAAAGAGGGAAGGGCCAAAGCTTTTTCTACCTGTGCATCCCACTTTTTGTCAGTTTCATTATTCTATGGATCTCTTTTCTTCATGTACGTTAGACCAAATTTGCTTGAAGAAGGGGATAAAGATATACCAGCTGCAATTTTATTTACAATAGTAGTTCCCTTACTAAATCCTTTCATTTATAGCCTGAGAAATAGGGAAGTAATAAGTGTCTTAAGAAAAATTCTGATGAAGAAATAA |
| **Restriction Site** | NheI/HindIII |
| **Primer sequence** | CTAACTTCTACGTGCCCTTCTCCAA (forward, TM=60℃, 5'→3')  ACTGCATTCTAGTTGTGGTTTGTCC (reverse, TM=60℃, 5'→3') |

**Table S1 Information for *Homo sapiens* OR5K1 and OR2W1**

**Table S1 continued**

| **Gene Name** | **Rho-OR2W1** |
| --- | --- |
| **Gene Sequence** | ACGCGTATGAACGGCACCGAGGGCCCAAACTTCTACGTGCCTTTCTCCAACAAGACAGGAGTGGTCATGGACCAAAGCAATTATAGTTCTTTACATGGTTTTATTCTGCTTGGCTTCTCTAACCATCCAAAAATGGAGATGATCCTGTCAGGAGTTGTCGCCATCTTCTACTTAATTACATTGGTGGGTAACACAGCCATCATTCTTGCATCTCTCCTGGATTCCCAGCTTCATACACCAATGTACTTTTTCCTCAGAAATTTATCTTTCCTAGATCTATGTTTCACAACCAGCATCATCCCTCAGGTGCTGGTCAACTTGTGGGGACCTGATAAGACCATCAGCTATGTGGGTTGTATCATCCAACTCTATGTTTACATGTGGTTGGGCTCAGTTGAGTGCCTTCTCCTGGCTGTTATGTCCTATGATCGTTTTACAGCTATATGTAAGCCCTTGCATTATTTTGTAGTCATGAACCCACATCTATGTCTAAAGATGATTATCATGATCTGGAGTATTAGTTTGGCCAATTCTGTAGTATTATGTACACTCACTCTGAATTTGCCCACATGTGGAAACAACATTCTGGATCATTTCTTGTGTGAGTTGCCAGCTCTGGTCAAGATAGCTTGTGTAGACACCACAACAGTTGAAATGTCTGTTTTCGCTTTAGGCATTATAATTGTCCTCACACCTCTCATCCTTATTCTTATATCCTATGGCTACATTGCCAAAGCTGTGCTGAGAACGAAGTCAAAAGCAAGCCAGCGAAAAGCAATGAATACCTGTGGATCTCATCTTACTGTAGTGTCTATGTTCTATGGAACTATTATCTACATGTACCTGCAACCAGGTAACAGGGCTTCCAAAGACCAGGGCAAGTTCCTCACCCTCTTTTACACTGTCATCACTCCAAGTCTCAACCCGCTCATTTACACCTTAAGAAATAAGGACATGAAGGATGCCCTGAAGAAACTGATGAGATTTCACCACAAATCTACAAAAATAAAGAGGAATTGCAAGTCATAG |
| **Restriction Site** | NheI/HindIII |
| **Primer sequence** | TAACTTCTACGTGCCCTTCTCCAAT (forward, TM=60℃, 5'→3')  GTATCTTATCATGTCTGCTCGAAGC (reverse, TM=58℃, 5'→3') |

**Table S2 Information for all ORs used for screening (excluding mutant variants)**

| **Num.** | **Gene Name** | **Haplotype** | **Mutant Status** |
| --- | --- | --- | --- |
| 1 | OR1A1 | OR1A1 | N. |
| 2 | OR1A2 | OR1A2 | N. |
| 3 | OR1B1 | OR1B1 | N. |
| 4 | OR1C1 | OR1C1 | N. |
| 5 | OR1D2 | OR1D2 | N. |
| 6 | OR1D5 | OR1D5 | N. |
| 7 | OR1E1 | OR1E1 | N. |
| 8 | OR1E2 | OR1E2 | N. |
| 9 | OR1E3 | OR1E3 | N. |
| 10 | OR1F1 | OR1F1 | N. |
| 11 | OR1G1 | OR1G1 | N. |
| 12 | OR1I1 | OR1I1 | N. |
| 13 | OR1J2 | OR1J2 | N. |
| 14 | OR1J4 | OR1J4 | N. |
| 15 | OR1K1 | OR1K1 | N. |
| 16 | OR1L1 | OR1L1 | N. |
| 17 | OR1L6 | OR1L6 | N. |
| 18 | OR1L8 | OR1L8 | N. |
| 19 | OR1M1 | OR1M1 | N. |
| 20 | OR1N1 | OR1N1 | N. |
| 21 | OR1N2 | OR1N2 | N. |
| 22 | OR1P1 | OR1P1 | N. |
| 23 | OR1Q1 | OR1Q1 | N. |
| 24 | OR1S1 | OR1S1 | N. |
| 25 | OR1S2 | OR1S2 | N. |
| 26 | OR2A1 | OR2A1 | N. |
| 27 | OR2A2 | OR2A2 | N. |
| 28 | OR2A4 | OR2A4 | N. |
| 29 | OR2A5 | OR2A5 | N. |
| 30 | OR2A7 | OR2A7 | N. |
| 31 | OR2A12 | OR2A12 | N. |
| 32 | OR2A14 | OR2A14 | N. |
| 33 | OR2A25 | OR2A25 | N. |
| 34 | OR2AE1 | OR2AE1 | N. |
| 35 | OR2AG2 | OR2AG2 | N. |
| 36 | OR2AJ1 | OR2AJ1 | N. |
| 37 | OR2AP1 | OR2AP1 | N. |
| 38 | OR2AT4 | OR2AT4 | N. |
| 39 | OR2B2 | OR2B2 | N. |
| 40 | OR2B3 | OR2B3 | N. |
| 41 | OR2B6 | OR2B6 | N. |
| 42 | OR2B8 | OR2B8 | N. |
| 43 | OR2B11 | OR2B11 | N. |
| 44 | OR2D3 | OR2D3 | N. |
| 45 | OR2F1 | OR2F1 | N. |
| 46 | OR2F2 | OR2F2 | N. |
| 47 | OR2G2 | OR2G2 | N. |
| 48 | OR2G3 | OR2G3 | N. |
| 49 | OR2G6 | OR2G6 | N. |
| 50 | OR2H1 | OR2H1 | N. |
| 51 | OR2H2 | OR2H2 | N. |
| 52 | OR2J2 | OR2J2 | N. |
| 53 | OR2J3 | OR2J3 | N. |
| 54 | OR2K2 | OR2K2 | N. |
| 55 | OR2L2 | OR2L2 | N. |
| 56 | OR2L8 | OR2L8 | N. |
| 57 | OR2L13 | OR2L13 | N. |
| 58 | OR2M2 | OR2M2 | N. |
| 59 | OR2M3 | OR2M3 | N. |
| 60 | OR2M4 | OR2M4 | N. |
| 61 | OR2M7 | OR2M7 | N. |
| 62 | OR2S2 | OR2S2 | N. |
| 63 | OR2T1 | OR2T1 | N. |
| 64 | OR2T2 | OR2T2 | N. |
| 65 | OR2T3 | OR2T3 | N. |
| 66 | OR2T4 | OR2T4 | N. |
| 67 | OR2T5 | OR2T5 | N. |
| 68 | OR2T7 | OR2T7 | N. |
| 69 | OR2T8 | OR2T8 | N. |
| 70 | OR2T10 | OR2T10 | N. |
| 71 | OR2T11 | OR2T11 | N. |
| 72 | OR2T27 | OR2T27 | N. |
| 73 | OR2T29 | OR2T29 | N. |
| 74 | OR2T34 | OR2T34 | N. |
| 75 | OR2V2 | OR2V2 | N. |
| 76 | OR2W1 | OR2W1 | N. |
| 77 | OR2W3 | OR2W3 | N. |
| 78 | OR2W5 | OR2W5 | N. |
| 79 | OR2Y1 | OR2Y1 | N. |
| 80 | OR3A1 | OR3A1 | N. |
| 81 | OR3A2 | OR3A2 | N. |
| 82 | OR3A3 | OR3A3 | N. |
| 83 | OR4A16 | OR4A16 | N. |
| 84 | OR4B1 | OR4B1 | N. |
| 85 | OR4C3 | OR4C3 | N. |
| 86 | OR4C5 | OR4C5 | N. |
| 87 | OR4C6 | OR4C6 | N. |
| 88 | OR4C13 | OR4C13 | N. |
| 89 | OR4C16 | OR4C16 | N. |
| 90 | OR4C45 | OR4C45 | N. |
| 91 | OR4C46 | OR4C46 | N. |
| 92 | OR4D1 | OR4D1 | N. |
| 93 | OR4D2 | OR4D2 | N. |
| 94 | OR4D5 | OR4D5 | N. |
| 95 | OR4D6 | OR4D6 | N. |
| 96 | OR4D9 | OR4D9 | N. |
| 97 | OR4D10 | OR4D10 | N. |
| 98 | OR4F3/16/29 | OR4F3/16/29 | N. |
| 99 | OR4F6 | OR4F6 | N. |
| 100 | OR4F14 | OR4F14 | N. |
| 101 | OR4F15 | OR4F15 | N. |
| 102 | OR4F17 | OR4F17 | N. |
| 103 | OR4F21 | OR4F21 | N. |
| 104 | OR4L1 | OR4L1 | N. |
| 105 | OR4K1 | OR4K1 | N. |
| 106 | OR4K2 | OR4K2 | N. |
| 107 | OR4K3 | OR4K3 | N. |
| 108 | OR4K14 | OR4K14 | N. |
| 109 | OR4K15 | OR4K15 | N. |
| 110 | OR4K17 | OR4K17 | N. |
| 111 | OR4M2 | OR4M2 | N. |
| 112 | OR4N2 | OR4N2 | N. |
| 113 | OR4P4 | OR4P4 | N. |
| 114 | OR4Q2 | OR4Q2 | N. |
| 115 | OR4S2 | OR4S2 | N. |
| 116 | OR4X1 | OR4X1 | N. |
| 117 | OR4X2 | OR4X2 | N. |
| 118 | OR5A1 | OR5A1 | N. |
| 119 | OR5A2 | OR5A2 | N. |
| 120 | OR5AC2 | OR5AC2 | N. |
| 121 | OR5AK3 | OR5AK3 | N. |
| 122 | OR5AL1 | OR5AL1 | N. |
| 123 | OR5AP2 | OR5AP2 | N. |
| 124 | OR5AS1 | OR5AS1 | N. |
| 125 | OR5AU1 | OR5AU1 | N. |
| 126 | OR5B2 | OR5B2 | N. |
| 127 | OR5B3 | OR5B3 | N. |
| 128 | OR5B12 | OR5B12 | N. |
| 129 | OR5B17 | OR5B17 | N. |
| 130 | OR5B21 | OR5B21 | N. |
| 131 | OR5C1 | OR5C1 | N. |
| 132 | OR5D13 | OR5D13 | N. |
| 133 | OR5D14 | OR5D14 | N. |
| 134 | OR5D16 | OR5D16 | N. |
| 135 | OR5F1 | OR5F1 | N. |
| 136 | OR5G3 | OR5G3 | N. |
| 137 | OR5H1 | OR5H1 | N. |
| 138 | OR5H2 | OR5H2 | N. |
| 139 | OR5H6 | OR5H6 | N. |
| 140 | OR5H14 | OR5H14 | N. |
| 141 | OR5I1 | OR5I1 | N. |
| 142 | OR5J2 | OR5J2 | N. |
| 143 | OR5K1 | OR5K1 | N. |
| 144 | OR5K2 | OR5K2 | N. |
| 145 | OR5K3 | OR5K3 | N. |
| 146 | OR5L1 | OR5L1 | N. |
| 147 | OR5M3 | OR5M3 | N. |
| 148 | OR5M8 | OR5M8 | N. |
| 149 | OR5M9 | OR5M9 | N. |
| 150 | OR5M11 | OR5M11 | N. |
| 151 | OR5P55 | OR5P55 | N. |
| 152 | OR5R1 | OR5R1 | N. |
| 153 | OR5S1 | OR5S1 | N. |
| 154 | OR5T2 | OR5T2 | N. |
| 155 | OR5T3 | OR5T3 | N. |
| 156 | OR5V1 | OR5V1 | N. |
| 157 | OR5W2 | OR5W2 | N. |
| 158 | OR6A2 | OR6A2 | N. |
| 159 | OR6B1 | OR6B1 | N. |
| 160 | OR6B2 | OR6B2 | N. |
| 161 | OR6B3 | OR6B3 | N. |
| 162 | OR6C1 | OR6C1 | N. |
| 163 | OR6C2 | OR6C2 | N. |
| 164 | OR6C3 | OR6C3 | N. |
| 165 | OR6C4 | OR6C4 | N. |
| 166 | OR6C65 | OR6C65 | N. |
| 167 | OR6C74 | OR6C74 | N. |
| 168 | OR6C75 | OR6C75 | N. |
| 169 | OR6C76 | OR6C76 | N. |
| 170 | OR6F1 | OR6F1 | N. |
| 171 | OR6K2 | OR6K2 | N. |
| 172 | OR6K3 | OR6K3 | N. |
| 173 | OR6K6 | OR6K6 | N. |
| 174 | OR6M1 | OR6M1 | N. |
| 175 | OR6N1 | OR6N1 | N. |
| 176 | OR6P1 | OR6P1 | N. |
| 177 | OR6S1 | OR6S1 | N. |
| 178 | OR7A5 | OR7A5 | N. |
| 179 | OR7A10 | OR7A10 | N. |
| 180 | OR7A17 | OR7A17 | N. |
| 181 | OR7D2 | OR7D2 | N. |
| 182 | OR7E24 | OR7E24 | N. |
| 183 | OR7G1 | OR7G1 | N. |
| 184 | OR8A1 | OR8A1 | N. |
| 185 | OR8B2 | OR8B2 | N. |
| 186 | OR8B4 | OR8B4 | N. |
| 187 | OR8B8 | OR8B8 | N. |
| 188 | OR8D1 | OR8D1 | N. |
| 189 | OR8D2 | OR8D2 | N. |
| 190 | OR8D4 | OR8D4 | N. |
| 191 | OR8G1 | OR8G1 | N. |
| 192 | OR8G5 | OR8G5 | N. |
| 193 | OR8H1 | OR8H1 | N. |
| 194 | OR8H2 | OR8H2 | N. |
| 195 | OR8H3 | OR8H3 | N. |
| 196 | OR8I2 | OR8I2 | N. |
| 197 | OR8J1 | OR8J1 | N. |
| 198 | OR8K1 | OR8K1 | N. |
| 199 | OR8K3 | OR8K3 | N. |
| 200 | OR8S1 | OR8S1 | N. |
| 201 | OR8U8 | OR8U8 | N. |
| 202 | OR8U9 | OR8U9 | N. |
| 203 | OR9A2 | OR9A2 | N. |
| 204 | OR9A4 | OR9A4 | N. |
| 205 | OR9G1 | OR9G1 | N. |
| 206 | OR9G9 | OR9G9 | N. |
| 207 | OR9I1 | OR9I1 | N. |
| 208 | OR9K2 | OR9K2 | N. |
| 209 | OR9Q2 | OR9Q2 | N. |
| 210 | OR10A3 | OR10A3 | N. |
| 211 | OR10A4 | OR10A4 | N. |
| 212 | OR10AD1 | OR10AD1 | N. |
| 213 | OR10AG1 | OR10AG1 | N. |
| 214 | OR10C1 | OR10C1 | N. |
| 215 | OR10D3 | OR10D3 | N. |
| 216 | OR10G3 | OR10G3 | N. |
| 217 | OR10G4 | OR10G4 | N. |
| 218 | OR10G8 | OR10G8 | N. |
| 219 | OR10G9 | OR10G9 | N. |
| 220 | OR10H2 | OR10H2 | N. |
| 221 | OR10H3 | OR10H3 | N. |
| 222 | OR10H4 | OR10H4 | N. |
| 223 | OR10J1 | OR10J1 | N. |
| 224 | OR10J5 | OR10J5 | N. |
| 225 | OR10K1 | OR10K1 | N. |
| 226 | OR10Q1 | OR10Q1 | N. |
| 227 | OR10R2 | OR10R2 | N. |
| 228 | OR10S1 | OR10S1 | N. |
| 229 | OR10T2 | OR10T2 | N. |
| 230 | OR10V1 | OR10V1 | N. |
| 231 | OR10W1 | OR10W1 | N. |
| 232 | OR10X1 | OR10X1 | N. |
| 233 | OR10Z1 | OR10Z1 | N. |
| 234 | OR11A1 | OR11A1 | N. |
| 235 | OR11G2 | OR11G2 | N. |
| 236 | OR11H2 | OR11H2 | N. |
| 237 | OR11H4 | OR11H4 | N. |
| 238 | OR11H6 | OR11H6 | N. |
| 239 | OR11H12 | OR11H12 | N. |
| 240 | OR11L1 | OR11L1 | N. |
| 241 | OR12D2 | OR12D2 | N. |
| 242 | OR12D3 | OR12D3 | N. |
| 243 | OR13C2 | OR13C2 | N. |
| 244 | OR13C3 | OR13C3 | N. |
| 245 | OR13C4 | OR13C4 | N. |
| 246 | OR13C5 | OR13C5 | N. |
| 247 | OR13C7 | OR13C7 | N. |
| 248 | OR13C8 | OR13C8 | N. |
| 249 | OR13C9 | OR13C9 | N. |
| 250 | OR13D1 | OR13D1 | N. |
| 251 | OR13F1 | OR13F1 | N. |
| 252 | OR13H1 | OR13H1 | N. |
| 253 | OR13J1 | OR13J1 | N. |
| 254 | OR14A2 | OR14A2 | N. |
| 255 | OR14C36 | OR14C36 | N. |
| 256 | OR14I1 | OR14I1 | N. |
| 257 | OR14J1 | OR14J1 | N. |
| 258 | OR14K1 | OR14K1 | N. |
| 259 | OR14L1 | OR14L1 | N. |
| 260 | OR51B2 | OR51B2 | N. |
| 261 | OR51B4 | OR51B4 | N. |
| 262 | OR51B5 | OR51B5 | N. |
| 263 | OR51B6 | OR51B6 | N. |
| 264 | OR51D1 | OR51D1 | N. |
| 265 | OR510 | OR510 | N. |
| 266 | OR51F1 | OR51F1 | N. |
| 267 | OR51G1 | OR51G1 | N. |
| 268 | OR51G2 | OR51G2 | N. |
| 269 | OR51H1 | OR51H1 | N. |
| 270 | OR51I2 | OR51I2 | N. |
| 271 | OR51L1 | OR51L1 | N. |
| 272 | OR51T1 | OR51T1 | N. |
| 273 | OR52A1 | OR52A1 | N. |
| 274 | OR52B4 | OR52B4 | N. |
| 275 | OR52B6 | OR52B6 | N. |
| 276 | OR52D1 | OR52D1 | N. |
| 277 | OR52E1 | OR52E1 | N. |
| 278 | OR52E4 | OR52E4 | N. |
| 279 | OR52E6 | OR52E6 | N. |
| 280 | OR52E8 | OR52E8 | N. |
| 281 | OR52I2 | OR52I2 | N. |
| 282 | OR52J3 | OR52J3 | N. |
| 283 | OR52K1 | OR52K1 | N. |
| 284 | OR52K2 | OR52K2 | N. |
| 285 | OR52L1 | OR52L1 | N. |
| 286 | OR52L2 | OR52L2 | N. |
| 287 | OR52N1 | OR52N1 | N. |
| 288 | OR52N4 | OR52N4 | N. |
| 289 | OR52N5 | OR52N5 | N. |
| 290 | OR52P1 | OR52P1 | N. |
| 291 | OR52W1 | OR52W1 | N. |
| 292 | OR56A3 | OR56A3 | N. |
| 293 | OR56B1 | OR56B1 | N. |
| 294 | OR56B4 | OR56B4 | N. |

**Table S3 Interpretation of units of measure**

| **Symbol** | **Full Name** | **Application Scenario** |
| --- | --- | --- |
| μM | Micromolar （μmol/L） | Concentration |
| g/L | grams per liter | Mass concentration |
| % | percent | Concentration/proportion |
| U/mL | units per milliliter | Bioactive substance concentration |
| μmol/L | micromoles per liter | Concentration (equivalent to μM) |
| M | molar | High-concentration solutions |
| cm | centimeter | Length |
| mL | milliliter | Volume |
| ng | nanogram | Mass |
| s | second | Time |
| kDa | kilodalton | Molecular weight |
| °C | degrees Celsius | Temperature |

**Table S4 Calculate the threshold value based on the S-curve method**

|  | ***x0*** | **Threshold (=10^*x0,* μM)** | ***b*** |
| --- | --- | --- | --- |
| **TMP** | 3.97197 | 9374.972447 | 0.32876 |
| **L-MT** | 4.12023 | 13189.550651 | 0.26578 |
| **TMP**+L-MT  (L-MT 100 mM） | 4.52216 | 33278.213218 | 0.40141 |
| **TMP+**L-MT(1:1） | 4.13505 | 13647.402491 | 0.41293 |
| **L-MT+**TMP  (TMP 1000 mM） | 4.72752 | 53397.386341 | 0.3135 |
| **L-MT+**TMP(1:1） | 4.24233 | 17471.492254 | 0.21089 |

*x0* and *b* are the parameters in the S-curve plot, derived from the function.

**Figure S1 Information for the vector of target gene plasmids**

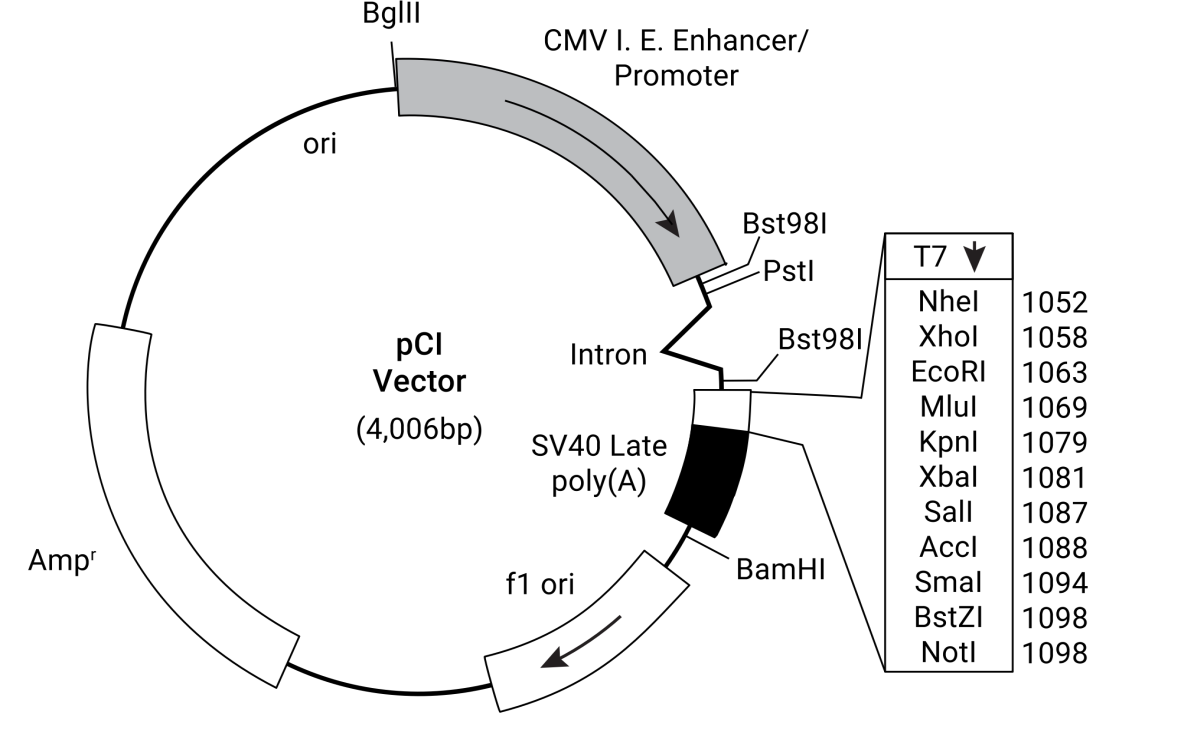

The pCI Mammalian Expression Vector promotes constitutive expression of cloned DNA inserts in mammalian cells. The major difference between the pCI and pSI Mammalian Expression Vectors is the enhancer/promoter region controlling expression of the inserted gene. The pCI Expression Vector contains the human cytomegalovirus (CMV) major immediate-early gene enhancer/promoter region. A β-globin/IgG chimeric intron located downstream of the enhancer/promoter region can further increase expression in the pCI Vector. The late SV40 polyadenylation signal increases the steady-state level of mRNA approximately fivefold more than the early SV40 polyadenylation signal.

(From: <https://www.promega.com.cn/)>

**Figure S2 A cellular experimental design protocol for exploring binary interactions of odorants**

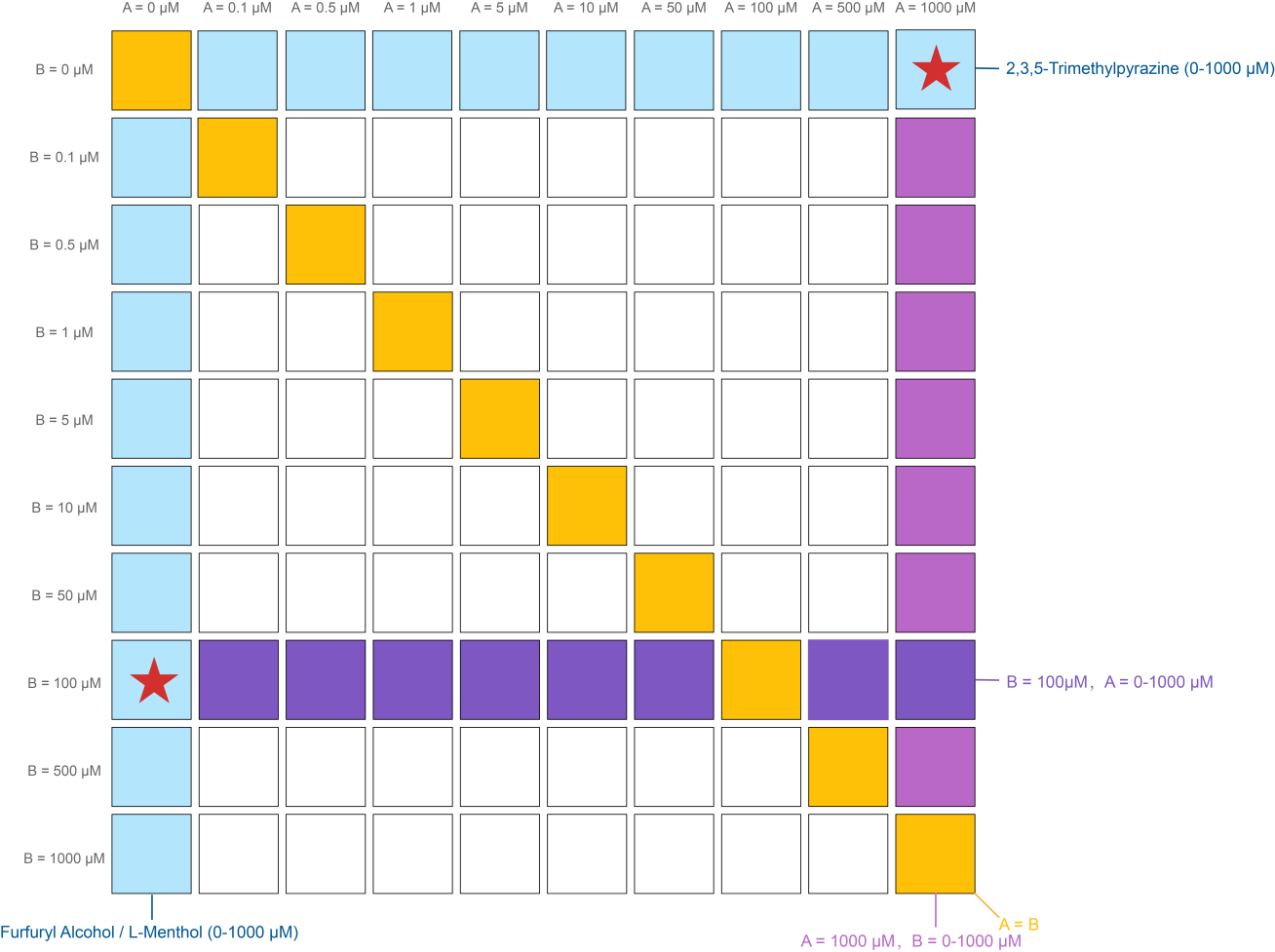

The experimental design employs a modular hierarchical architecture to integrate high-throughput screening with in-depth mechanistic analysis, focusing on TMP (A) and L-MT (B) interactions at olfactory receptor (OR) level. This innovative approach offers three key advantages: (1) Independent analysis of single-variable dose responses (A/B gradient: 0-1000 μM) identifies optimal concentrations (A10=1000 μM, purple squares; B8=100 μM, dark blue squares); (2) Diagonal experiments simplify symmetric interaction analysis, revealing synergistic/masking patterns in equimolar binary systems; (3) Dynamic expansion layers (A9 column/B8 row) systematically validate interaction modes between optimal and suboptimal concentrations. The core strength of this approach lies in its implementation of a “observation-verification-optimization” closed-loop research workflow through dynamic expansion interfaces. This design not only aligns with the “Stepwise Optimization” engineering logic but also incorporates modular architecture to accommodate future technical iterations, significantly enhancing both the efficiency and decision-making reliability of olfactory receptor research.

**Figure S3 Composition testing of binary systems**

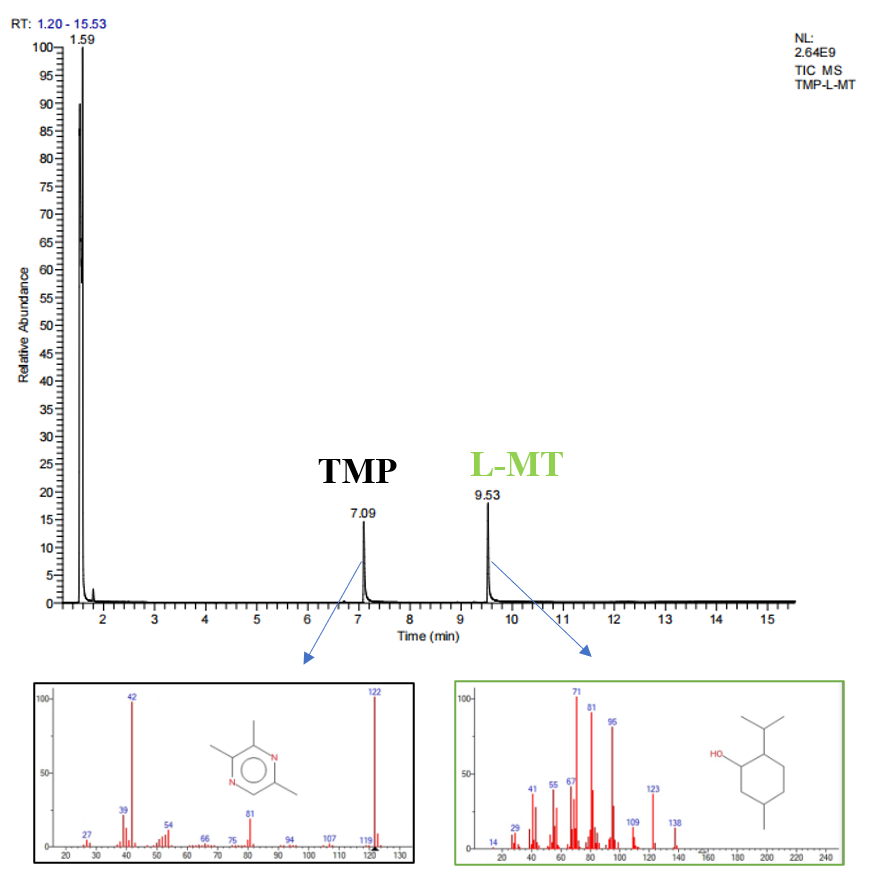

We employed gas chromatography-quadrupole/orbitrap mass spectrometry (GC-Q/Orbitrap MS, Thermo Fisher Scientific, America) to analyze pre-configured binary systems, revealing that no chemical reactions occurred between the components, and no new substances were formed. Each component in the binary systems remained stable and existed independently.

The separation was performed on a DB-5MS column (30 m × 0.25 mm × 0.25 µm) with ultra-pure helium (99.999%) as the carrier gas at a constant flow rate of 1.0 mL/min. The injector temperature was set to 250 °C with a split ratio of 50:1, and the injection volume was 1 µL. The oven temperature program started at 60 °C, held for 4 minutes, then increased to 240 °C at a rate of 15 °C/min, and held for 2 minutes.

The mass spectrometer was equipped with an electron ionization (EI) source operating at 70 eV. The ion source temperature was set to 250 °C, and the MS transfer line temperature was maintained at 290 °C. The mass range was set to m/z 35–525, and the Orbitrap resolution was 60,000.

This work was completed with the assistance of Professor Liu Yumin from the Analysis and Testing Center of Shanghai Jiao Tong University.

**Detailed instructions for sensory experiments**

Considering the discrepancy between human olfactory sensitivity and OR detection thresholds, coupled with volatile compound diffusion from solution to air phase, all concentrations employed in sensory experiments were systematically amplified by three orders of magnitude (1000×, from μM to mM) compared to those used in cellular assays. The test samples are randomly coded with three digits in an AAB, ABA and BAA manner to balance the order of provision of the three samples in front of the evaluator and avoid positional bias. Liquid samples should not exceed 1/4 of the volume of the container. Containers should be used that are stoppered and suitable for sniffing.

*Sample Preparation:*

In this study, five groups of test samples were set up and formulated as follows:

Group A: TMP solution with concentrations of 0.1 mM, 0.5 mM, 1 mM, 5 mM, 10 mM, 50 mM, 100 mM, 500 mM, and 1000 mM were prepared, respectively.

Group B: L-MT solution with concentrations of 0.1 mM, 0.5 mM, 1 mM, 5 mM, 10 mM, 50 mM, 100 mM, 500 mM, and 1000 mM were prepared, respectively.

Group C: equal concentrations of TMP and L-MT, divided into 0.1 mM, 0.5 mM, 1 mM, 5 mM, 10 mM, 50 mM, 100 mM, 500 mM, and 1000 mM;

Group D: the concentration of L-MT was fixed at 100 mM, the concentration of TMP was set at 0.1 mM, 0.5 mM, 1 mM, 5 mM, 10 mM, 50 mM, 100 mM, 500 mM, and 1000 mM, and the reference sample was TMP;

Group E: the concentration of TMP was fixed at 1000 mM, the concentration of L-MT was set at 0.1 mM, 0.5 mM, 1 mM, 5 mM, 10 mM, 50 mM, 100 mM, 500 mM, and 1000 mM, and the reference sample was L-MT;

All solutes for the sensory experiments were prepared stock with DMSO and then diluted with deionized water to the desired concentration.

*Sample Testing:*

In the 3-AFC tests, each target sample was presented alongside two solvent-only blanks. Panelists were required to identify the different sample among the triad. Threshold concentrations were calculated through statistical analysis of correct identification rates across concentration series.

- Groups A & B: Evaluators assessed individual odorant (TMP or L-MT) at varying concentrations.
- Group C: Participants performed dual evaluations of the same samples: First assessment is TMP detection; Second assessment is L-MT detection.
- Group D: Threshold determination for TMP in the presence of fixed-concentration L-MT background.
- Group E: Threshold determination for L-MT in the presence of fixed-concentration TMP background.
